# Macrophages stimulate epicardial VEGFaa expression to trigger cardiomyocyte proliferation in larval zebrafish heart regeneration

**DOI:** 10.1101/2021.06.15.448575

**Authors:** Finnius A. Bruton, Aryan Kaveh, Katherine M. Ross-Stewart, Gianfranco Matrone, Magdalena E.M. Oremek, Emmanouil G. Solomonidis, Carl S. Tucker, John J. Mullins, Mairi Brittan, Jonathan M. Taylor, Adriano G. Rossi, Martin A. Denvir

## Abstract

Cardiac injury induces a sustained macrophage response in both zebrafish and mammals. Macrophages perform a range of both beneficial and detrimental functions during mammalian cardiac repair, yet their precise roles in zebrafish cardiac regeneration are not fully understood. Here we characterise cardiac regeneration in the rapidly regenerating larval zebrafish laser injury model and use macrophage ablation and macrophage-null *irf8* mutants to define the role of macrophages in key stages of regeneration. Live heartbeat-synchronised imaging and RNA sequencing revealed an early proinflammatory phase, marked by tnfa+ macrophages, which then resolved to an anti-inflammatory, profibrotic phase. Macrophages were required for cardiomyocyte proliferation but not for functional or structural recovery following injury. Importantly, we found that macrophages are specifically recruited to the epicardial-myocardial niche, triggering the expansion of the epicardium which upregulates VEGFaa expression to induce cardiomyocyte proliferation. Hence, revealing a novel mechanism by which macrophages facilitate cardiac regeneration.

## Introduction

Zebrafish are highly regenerative, exhibiting the capacity to restore full structure and function to a wide range of tissues following injury^1–5^. Cardiac injury is one such example where adult mammals are only able to facilitate maladaptive repair but zebrafish exhibit full tissue regeneration^6, 7^. In humans, the most severe form of cardiac injury is myocardial infarction (MI), where occlusion of a coronary artery triggers ischemic injury to the myocardium, leading to the loss of approximately 1 billion cardiomyocytes^8^. Adult mammalian cardiomyocytes are considered largely post-mitotic, switching to hypertrophic growth shortly after birth. They are therefore unable to restore lost myocardium, which is instead replaced with non-contractile scar tissue^9^. Consequently, MI patients suffer sequalae of maladaptive remodelling, leading to left ventricular dilation and thinning of the scar, further decreasing the function of the heart^10, 11^. Hence, there is a need for medical innovation which can reverse or prevent this process.

In contrast to mammalian models of MI, apical resection and cryoinjury MI models in zebrafish show full regeneration of lost myocardium *via* the dedifferentiation and proliferation of surviving cardiomyocytes^12, 13^. Cardiac regeneration is complex and dynamic, with zebrafish hearts undergoing debridement of dead myocardium, followed by transient fibrosis, revascularisation and eventual replacement of cardiomyocytes^14^. The inflammatory response has been demonstrated to be crucial for each of these key events, both in zebrafish and also in other regenerative species such as axolotls and neonatal mice^15–17^. In particular, macrophages have emerged as important cellular regulators of tissue regeneration. Indeed, macrophage ablation has been shown to abrogate regeneration across multiple organs and organisms, including the adult zebrafish heart^15, 16, 18^. However, the precise contribution of macrophages to cardiac repair has been complicated by disparate results following macrophage perturbation in mouse models of MI, where macrophages have been reported to be both beneficial and detrimental^7, 19, 20^. This is in part attributed to substantial heterogeneity of macrophage subtypes, and phenotypic plasticity^21, 22^. Recent studies have confirmed the presence of macrophage subsets in zebrafish, yet their functional niche and interactions with other key cell types of the heart, such as the epicardium, remain poorly understood^23, 24^. The larval zebrafish model of cardiac regeneration offers a tractable system to examine macrophages in detail. Larval zebrafish regenerate more rapidly than adults, occurring in just 48 hours after cardiac laser injury in 3-day old larvae^25, 26^. Combined with their amenability for live *in vivo* imaging and genetic tractability, this model becomes a powerful tool with which to carefully examine how macrophages support multiple aspects of cardiac regeneration.

Here we report an in-depth characterisation of the macrophage response and several key regenerative processes in larval zebrafish cardiac regeneration, finding the heart regeneration program between the larvae and adults to be highly conserved. Abolition of the macrophage response using metronidazole-nitroreductase ablation of macrophages or the macrophage null *irf8^-/-^* mutant^27^, demonstrated a requirement for these cells in removal of apoptotic cells, epicardial activation and cardiomyocyte proliferation. Interestingly, we found that one of the ways macrophages exert their pro-proliferative effect is via epicardial VEGFaa and downstream endocardial notch signalling. Our study reveals that macrophages invade the epicardial-myocardial niche, inducing expansion of epicardial cell numbers which increases epicardial VEGFaa expression, leading to an upregulation of endocardial notch signalling and the cardiac developmental growth pathway.

## Results

### Macrophages display cellular heterogeneity following cardiac injury

We first assessed macrophage heterogeneity and recruitment dynamics following larval cardiac injury. We crossed the zebrafish pan-macrophage reporter line *Tg(mpeg1:GFP)* with *Tg(csf1ra:gal4;UAS:mCherry-NfsB)* (shortened here to csf1ra: mCherry) (Supplementary Figure 1a). Csf1ra (colony stimulated factor 1 receptor) is a cytokine required for macrophage development and used as a macrophage reporter promoter in mammals^28^.

Larval hearts were lasered at the ventricular apex at 72 hours post-fertilisation (hpf) and imaged at 2, 6, 24, and 48 hours post injury (hpi) (Figure 1a). Macrophages migrate to the injured ventricular apex within 2 hours, peak at 6 and maintain elevated numbers until 48 hpi (Figures 1b & 1c). We found that not all recruited macrophages were co-positive for both transgenes, leading to three subsets 1) mpeg1+csf1ra-(19.3±5.1%), 2) mpeg1-csf1ra+ (2.8±2.1%) and 3) mpeg1+csf1ra+ (77.9±5.7%). Similar dynamics were seen for subsets 1 & 3 but since mpeg1-csf1ra+ were exceedingly rare it is not possible to know if the dynamics are likewise similar. Both subsets exhibit a range of morphologies with no overt difference between groups (Figure 1d, Video 1). Importantly, our data demonstrate that larval macrophages recruited to cardiac injury are heterogenous in their marker expression, similar to adult zebrafish^29^, and suggest a comparatively complex macrophage response in the larval model.

**Figure 1:**
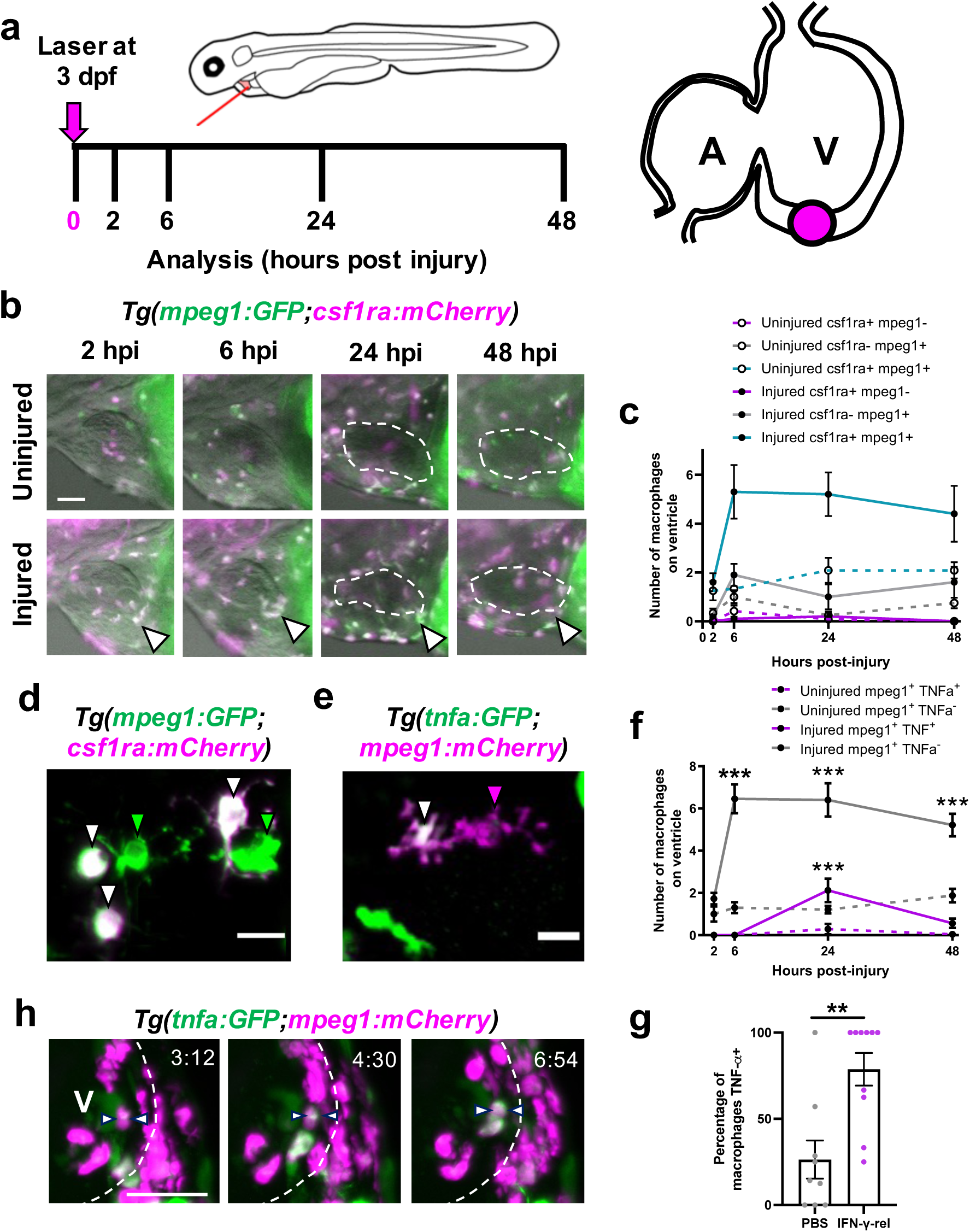
Cardiac macrophages display heterogeneity and plasticity following injury. (a) Schematic illustrating the cardiac laser injury model, with imaging timepoints marked (left) and the injury site at ventricular apex of a 3 dpf larval heart marked (magenta circle) (right). (b) Representative lateral view epifluorescence images of uninjured and injured hearts at the standard timepoints in *Tg(mpeg1:GFP;csf1ra:gal4:UAS:NfsB-mCherry)* (abbreviated to mpeg1:GFP;csf1ra:mCherry in all panels) illustrating macrophage heterogeneity, white arrow = ventricular apex; dashed line = heart outline. (c) Quantification of the number of csf1ra+mpeg1-, csf1ra-mpeg1+ and csf1ra+mpeg1+ macrophages on the ventricle in uninjured and injured larvae at standard timepoints, n=10-12. (d) Representative LSFM image of csf1ra-mpeg1+ and csf1ra+mpeg1+ macrophages of different morphologies. (e) Representative LSFM image of tnfa*+*mpeg1+ and tnfa*-* mpeg1+ macrophages. (f) Quantification of number of tnfa*+*mpeg1+ and tnfa*-*mpeg1+ macrophages on the ventricle in uninjured and injured larvae at standard timepoints, n=10-25. (g) Time-lapse timepoints for injured *Tg(tnfa:GFP;mpeg1:mCherry)* ventricles imaged live in the larvae by heartbeat-synchronised LSFM microscopy illustrating macrophage plasticity. Timestamps indicated, dashed line = ventricle outline; arrows = macrophage converting to tnfa*+.* (h) Quantification of the percentage of tnfa+ macrophages at 24 hpi following injection with IFN-γ-rel or PBS, n=10. Scale bar = 50μm (b & h), 10μm (d & e). *\*\*p≤0.01, ***p≤0.001*, (c & f) 2way ANOVA followed by Holm-Sidak’s Post-hoc test and (g) ttest.

### Macrophages display cellular plasticity following cardiac injury

To examine if macrophages display plasticity and convert to an inflammatory phenotype in the larval cardiac injury model, we performed cardiac laser injury on *Tg(tnf*a*:GFP;mpeg1:mCherry)* larvae. (Figure 1e & 1f). Quantification of tnfa+ macrophage number revealed a transient tnfa+ subset (19.3±4.9% of mpeg1+ macrophages, n=24), found only at the 24 hpi timepoint and rarely in uninjured larvae (Figure 1f & Supplementary figure 1b). We also observed that from 24 hpi, macrophages retract their pseudopods and become spherical, further suggesting a shift in phenotype (Supplementary figure 1e).

We reasoned that if tnfa+ macrophages were indeed inflammatory macrophages then application of M1-polarisation cytokine IFN-γ would increase their abundance. A single intravenous injection of zebrafish recombinant protein IFN-γ-rel, immediately prior to cardiac injury, increased the proportion of tnfα+mpeg1+ macrophages from 26.4±11.0% in PBS injected controls to 78.8±9.5%, supporting the suggestion that these were inflammatory macrophages (Figure 1g & Supplementary figure 1c & 1d).

Furthermore, *in vivo* imaging live in the beating heart showed recruited macrophages becoming tnfa:GFP+ after arrival at the injured ventricle, confirming that this represents true *in situ* conversion (Figure 1h, Video 2). Taken together, these data show that, as in adults, macrophages display plasticity and become inflammatory in response to cardiac injury, confirming the complexity of the macrophage response in this model.

### Larval cardiac laser lesions are similar in structure to adult cryoinjury

To validate analyses of macrophage function in the larval injury model, we first sought to determine if the laser lesion is comparable to adult cryoinjury and mammalian infarcts. Using the line *Tg(myl7:mKateCAAX;myl7:h2b-GFP)*, which labels cardiomyocyte sarcolemma and chromatin respectively, we observed that, following injury, a circlet of cardiomyocytes with pyknotic nuclei formed (Figure 2a). These pyknotic nuclei were TUNEL+ at 6 hpi, confirming apoptosis and they encircled the GFP- lesion (Figure 2b & 2c). Heartbeat-synchronised LSFM (lightsheet fluorescence microscopy)^30^ showed that nuclear condensation occurred extremely rapidly, being identified by 1.5 hpi (Supplementary figure 2a, Video 3).

**Figure 2:**
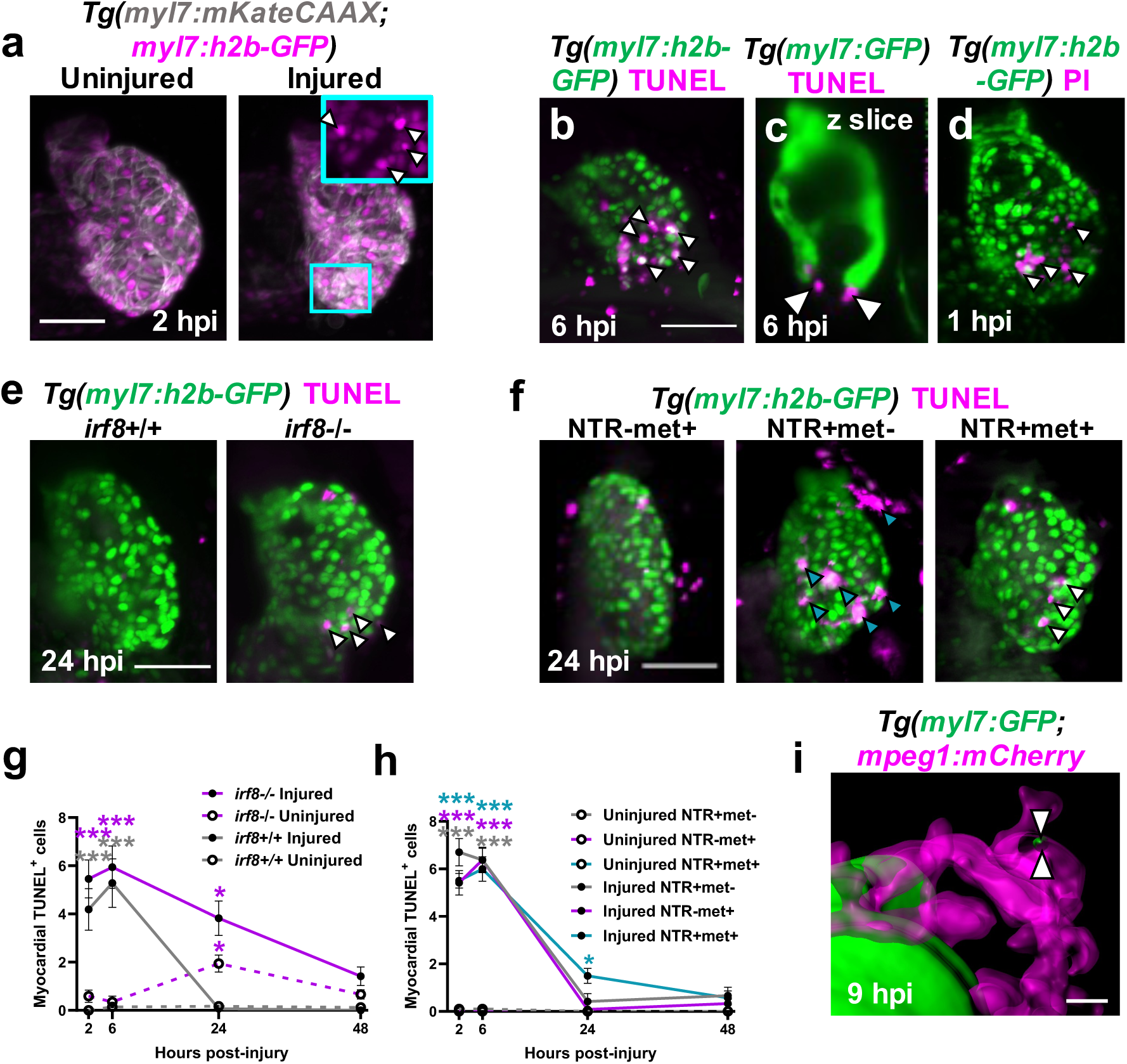
Macrophages are required for timely removal of apoptotic cardiomyocytes. (a) Representative LSFM images of uninjured and injured *Tg(myl7:h2b-GFP;myl7:mKateCAAX)* ventricles. Cyan outlined zoom panel highlights condensed nuclei (white arrowheads). Representative LSFM images of TUNEL stained hearts 6 hpi in (b) *Tg(myl7:h2b-GFP)* and (c) *Tg(myl7:GFP)* larvae. White arrowheads = apoptotic cardiomyocytes/myocardium. (d) Representative LSFM image of a propidium iodide (PI) stained *Tg(myl7:h2b-GFP)* heart at 1 hpi. White arrowheads = necrotic debris. (e) Representative LSFM images of *irf8^+/+^* and *irf8^-/-^ Tg(myl7:h2b-GFP)* hearts stained by TUNEL at 24 hpi. White arrowheads = TUNEL+ cells. (f) Representative LSFM images of injured *Tg(myl7:h2b-GFP;csfr1a:NfsB-mCherry)* ventricles per macrophage ablation model injury group at 24 hpi. Cyan arrowheads = Macrophages and white arrowheads = TUNEL+ cells. (g) Quantification of TUNEL+myocardial cells in uninjured and injured, *irf8^+/+^* and *irf8^-/-^ Tg(myl7:h2b-GFP)* ventricles, n=15-29. (h) Quantification of TUNEL+ myocardial cells in uninjured and injured *Tg(myl7:h2b-GFP;csfr1a:NfsB-mCherry)* ventricles per macrophage ablation group, n=10-12. (i) Surface render of LSFM-acquired z stack, surfaced-rendered with 50 % transparency at 9 hpi, showing internalised myocardial debris (white arrowheads) in a macrophage in a *Tg(myl7:GFP;mpeg1:mCherry)* larva. Scale bars = 50 μm for panels a-f and 20 μm for panel (i). All images are 3D LSFM shown as maximum intensity projections unless otherwise stated. \**p≤0.05,* \*\**p≤0.01,* \*\*\**p≤0.001* 2way ANOVA followed by Holm-Sidak’s Post-hoc tests.

We hypothesised that the GFP- epicentre of the laser lesion may contain cells that immediately necrose upon injury. To test this, we labelled necrotic cells by injecting propidium iodide (PI) intravenously immediately following injury (<0.5 hpi). We found that there were indeed PI+ cells in the GFP- region, and PI+ debris scattered across the proximal myocardium from 1 hpi (Figure 2d). Time-lapse imaging of PI-injected, injured *Tg(mpeg1:GFP;myl7:h2b-GFP;myl7:mKateCAAX)* hearts showed that necrotic cells are rapidly cleared within the first 0-2 hpi (Supplementary figure 2b & Video 3). Necrotic cells either disintegrated or were squeezed out from the myocardium into the pericardial cavity, independently of macrophage contact (Video 4). Overall, this characterisation confirms the structure of the laser lesion mirrors the necrotic infarct and apoptotic border zone observed in adult zebrafish cryoinjury and mammalian MI^14, 31^.

### Macrophages contribute to the removal of apoptotic cardiomyocytes following injury

We next sought to understand what role macrophages play in the regeneration of the larval heart, which occurs within only 48 hours of the initial injury^25, 26^. We used two different methods to induce macrophage-less hearts. Firstly we used the *Tg(csf1ra:gal4;UAS:mCherry-NfsB)* line (abbreviated hereafter to csf1ra:NfsB-mCherry) that expresses a nitroreductase enzyme NfsB in macrophages, which induces cell-specific apoptosis when exposed to prodrug metronidazole^32^ (Supplementary figure 3a-d). Macrophage ablation only occurs in larvae expressing the nitroreductase (NTR) *and* in the presence of metronidazole (NTR+met+). Therefore, larvae only expressing the nitroreductase (NTR+met-) or only in the presence of metronidazole (NTR-met+) are used as macrophage-replete control groups. The second method was the use of the macrophage-null *irf8*^-/-^ mutant^27^, IRF8 being a transcription factor required for macrophage development (Supplementary figure 3e-h).

To determine if macrophages are required for the removal of apoptotic cells, we performed TUNEL staining on *irf8*^-/-^ and *irf8^+/+^ Tg(myl7:h2b-GFP)* larvae at the standard 2, 6, 24, 48 hpi timepoints (Figure 2e & 2g). In injured *irf8*^+/+^ hearts, the number of apoptotic cardiomyocytes significantly increased at 2 hpi and 6 hpi compared to uninjured controls (4.1±0.9 vs 0.0±0.0 and 5.3±1.0 vs 0.1±0.1 respectively, n=15-29) but returned to baseline by 24 hpi. However, although injured macrophage-null *irf8*^-/-^ hearts showed a similar initial pattern of cell death at 2 hpi and 6 hpi (5.5±0.8 & 5.9±0.9 apoptotic cardiomyocytes respectively), apoptotic cardiomyocyte cells were still present at 24 hpi, only returning to uninjured levels by 48 hpi.

In the macrophage ablation model, we saw a similar pattern of results where the numbers of apoptotic cells were negligible in uninjured hearts of all treatment groups, but peaked at 6 hpi following injury (NTR+met-, NTR-met+ & NTR+met+ =6.4±0.5, 6.4±0.5 and 6.0±0.5, n=10-12) (Figure 2f & 2h). By 24 hpi the non-ablated groups no longer possessed significantly increased numbers of TUNEL+ myocardial cells; however, the macrophage-ablated group showed a retention of apoptotic cells at 24 hpi (NTR+met+ = 1.5±0.3) that resolved by 48 hpi.

To verify that macrophages are directly removing myocardial debris, we performed time-lapse imaging of injured *Tg(myl7:GFP;mpeg1:mCherry)* larvae. We observed small GFP+ pieces of myocardial debris near the GFP- lesion being removed and internalised by macrophages (Figure 2i & Supplementary Figure 2c & Video 5), confirming the essential role of macrophages in lesion debridement.

### Macrophages are not obligatory for structural or functional recovery of the larval heart

Next, we sought to investigate if macrophages are required for structural and functional recovery of the larval heart following laser-injury. We injured *Tg(myl7:GFP)* larvae following macrophage ablation and acquired serial 3D scans of the cardiac structure of individual larvae by heartbeat-synchronised LSFM (Figure 3a). In all treatment groups, the lesion size was consistent between 2 hpi and 6 hpi, with no difference between groups. By 24 hpi the lesion had almost completely regressed (95% to 37.1µm^2^±24.4, n=11-22) in macrophage-replete NTR+met- larvae (Figure 3b). However, for larvae in the macrophage-ablated NTR+met+ and the other macrophage-replete NTR-met+ group, lesion closure was slightly delayed at 24 hpi (73% and 75% to 234.7±59.7 µm^2^ and 221.4±84.6 µm^2^ respectively). By 48 hpi the lesions of larvae from each group had entirely regressed and luminal surface renders of injured ventricles showed normal trabecular structure (Figure 3a). These results suggest macrophages are not required for lesion closure, but that metronidazole-treatment slightly delays this process.

**Figure 3:**
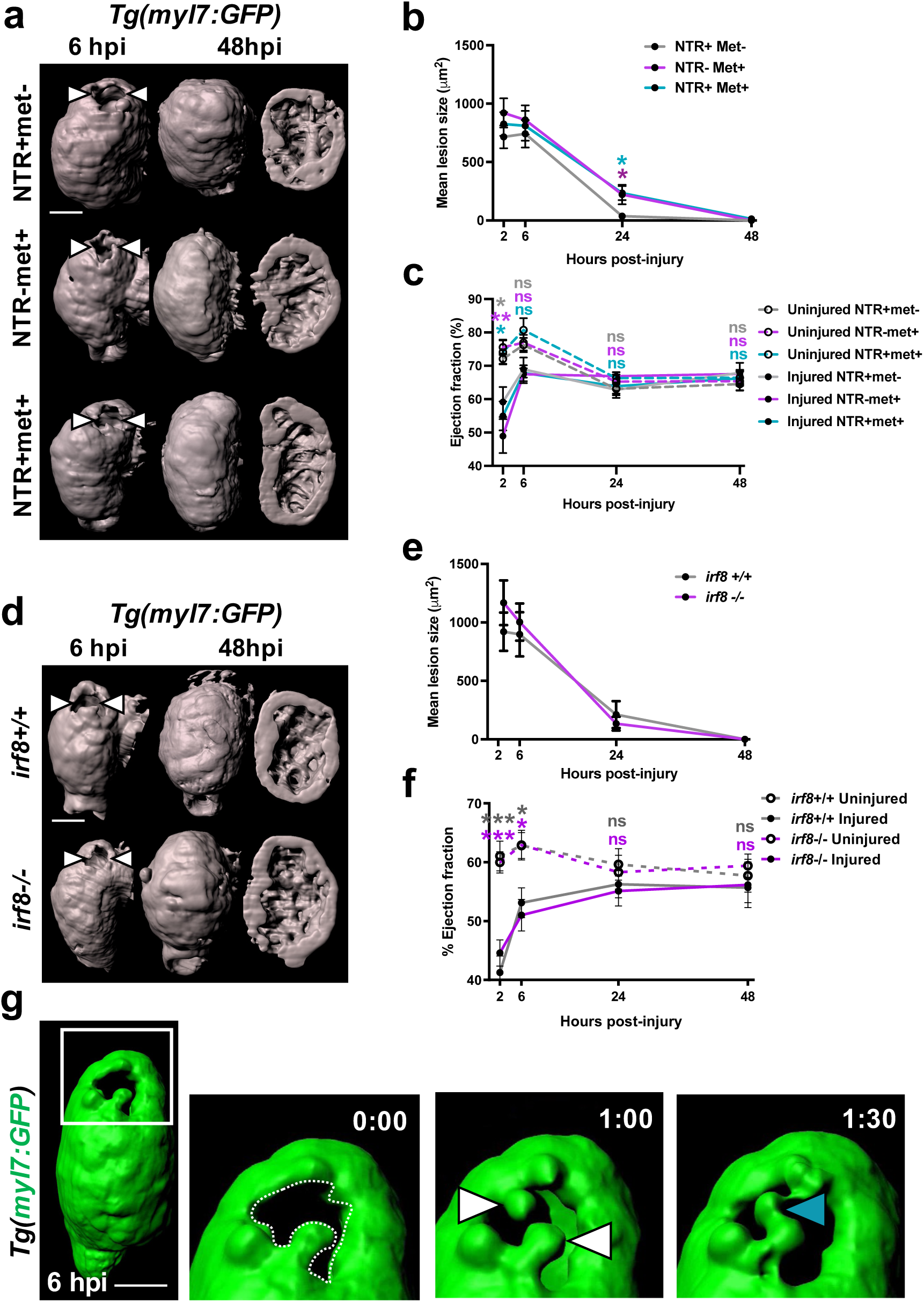
Macrophages not required for the recovery of cardiac structure or function. (a) Representative GFP surface-renders of LSFM z-stacks of injured ventricles in *Tg(myl7:GFP;csfr1a:NfsB-mCherry)* larvae, macrophage ablation groups as indicated in the figure. Abluminal myocardial surface is shown at 6 hpi (left) and abluminal and luminal surfaces shown at 48 hpi following regeneration (middle & right). White arrowheads = laser lesion. (b) Quantification of mean lesion size in injured *Tg(myl7:GFP;csfr1a:NfsB-mCherry)* larvae per macrophage ablation group, n=11-22. (c) Quantification of ventricular ejection fraction in uninjured and injured *Tg(myl7:GFP;csfr1a:NfsB-mCherry)* larvae per macrophage ablation group, n=10-12. (d) Representative GFP surface-renders of light-sheet-acquired z-stacks of injured ventricles from *irf8^+/+^* and *irf8^-/-^ Tg(myl7:GFP)* larvae. Abluminal myocardial surface is shown at 6 hpi (left) and abluminal and luminal surfaces shown at 48 hpi following regeneration (middle & right). White arrowheads = laser lesion. (e) Quantification of mean lesion size in injured *irf8^+/+^* and *irf8^-/-^ Tg(myl7:GFP)* larvae, n=15. (f) Quantification of ventricular ejection fraction in uninjured and injured *irf8^+/+^* and *irf8^-/-^ Tg(myl7:GFP)* larvae n=15-20. (g) Time-lapse timepoints of a GFP-surface-rendered, injured *Tg(myl7:GFP)* ventricle from 6 hpi. White box = zoom panel; white arrowheads = myocardial buds, cyan arrowhead = myocardial bridge. \**p≤0.05,* \*\*\**p≤0.001* 2way ANOVA followed by Holm-Sidak’s Post-hoc tests. Scale bars = 50 μm

Using *Tg(myl7:GFP)* larvae, we acquired lateral-view videos of beating hearts with epifluorescence microscopy and tested if macrophage ablation affected recovery of cardiac function. Immediately following injury at 2 hpi, volumetric ejection fraction was decreased in all groups from 74% in uninjured ventricles to 54% in injured (Figure 3c & Supplementary figure 4a & 4b). Ejection fraction recovered quickly by 6 hpi in all treatment groups, (∼78% injured vs ∼87% uninjured) and by 24 hpi and 48 hpi injured hearts were functionally indistinguishable from uninjured hearts. These data suggest that injured larval hearts recover their function rapidly, and that this recovery is not macrophage dependent.

We next performed identical experiments examining the recovery of cardiac structure and function with *Tg(myl7:GFP)* larvae on an *irf8* mutant background. Both *irf8*^+/+^ and *irf8*^-/-^ genotype larvae showed substantial lesion regression (∼80%) between 6 hpi and 24 hpi (898.6 µm^2^±189.7 to 211.6 µm^2^±115.8 vs 1002.9 µm^2^±158.4 to 113.89 µm^2^±59.5, respectively) (Figure 3d & 3e). No difference in lesion size was seen at any timepoint and both genotypes had completely closed their lesions by 48 hpi. Normal trabecular structure was seen in both groups at 48 hpi following full structural recovery (Figure 3d). The recovery of ejection fraction in this model followed the same trend as that of the metronidazole-nitroreductase model, with the ejection fraction of injured larvae being indistinguishable from uninjured larvae by 24 hpi in both genotypes (Figure 3f). Our near identical findings in the *irf8* macrophage null model confirm that larval hearts rapidly recover following laser injury and that this process is macrophage independent.

Finally, we wished to understand the mechanism of lesion closure. We performed heartbeat-synchronised time-lapse imaging of lesions in *Tg(myl7:GFP)* larvae immediately following injury. We observed GFP+ myocardial budding on opposite sides of the lesion border zone and subsequent invasion into the lesion, adhering to each other to form bridges (Figure 3g, Video 6). Repeating this experiment in *Tg(myl7:h2b-GFP;myl7:mKateCAAX)* larvae facilitated the tracking of individual cardiomyocytes by virtue of their labelled nuclei and plasma membranes (Video 7 & Supplementary figure 2d). We found that cardiomyocytes bordering the lesion did not divide but extended protrusions into the lesion until they adhered with other single cardiomyocytes bridging from the opposing side of the lesion. These imaging insights suggest that myocardial structure is first restored by morphogenesis rather than cell division.

### Macrophage ablation abolishes an injury-associated increase in cardiomyocyte proliferation

To test if cardiomyocyte proliferation increases in response to laser-injury, we performed EdU staining in *Tg(myl7:h2b-GFP)* larvae in two experiments. In the first experiment, uninjured and injured larvae were exposed to EdU during 0-24 hpi and then at 24-48 hpi for the second (Figure 4a). Comparison between uninjured and injured hearts revealed no significant difference in the proportion of EdU+ cardiomyocyte nuclei 0-24 hpi (21.3±3.3 vs 18.9±3.4 respectively, n=10-14) (Figure 4b & 4c). However, over 24-48 hpi there was an organ-wide, 35% increase in the proportion of EdU+ cardiomyocytes in injured hearts relative to uninjured (43.5±1.8% vs 32.2±2.0% respectively, n=17-25). Time-lapse *in vivo* imaging of dividing cardiomyocytes showed nuclear division followed by cytokinesis, exclusively gives rise to mononuclear cells, with no obvious hypertrophy (Video 8, Supplementary Figure 5a).

**Figure 4:**
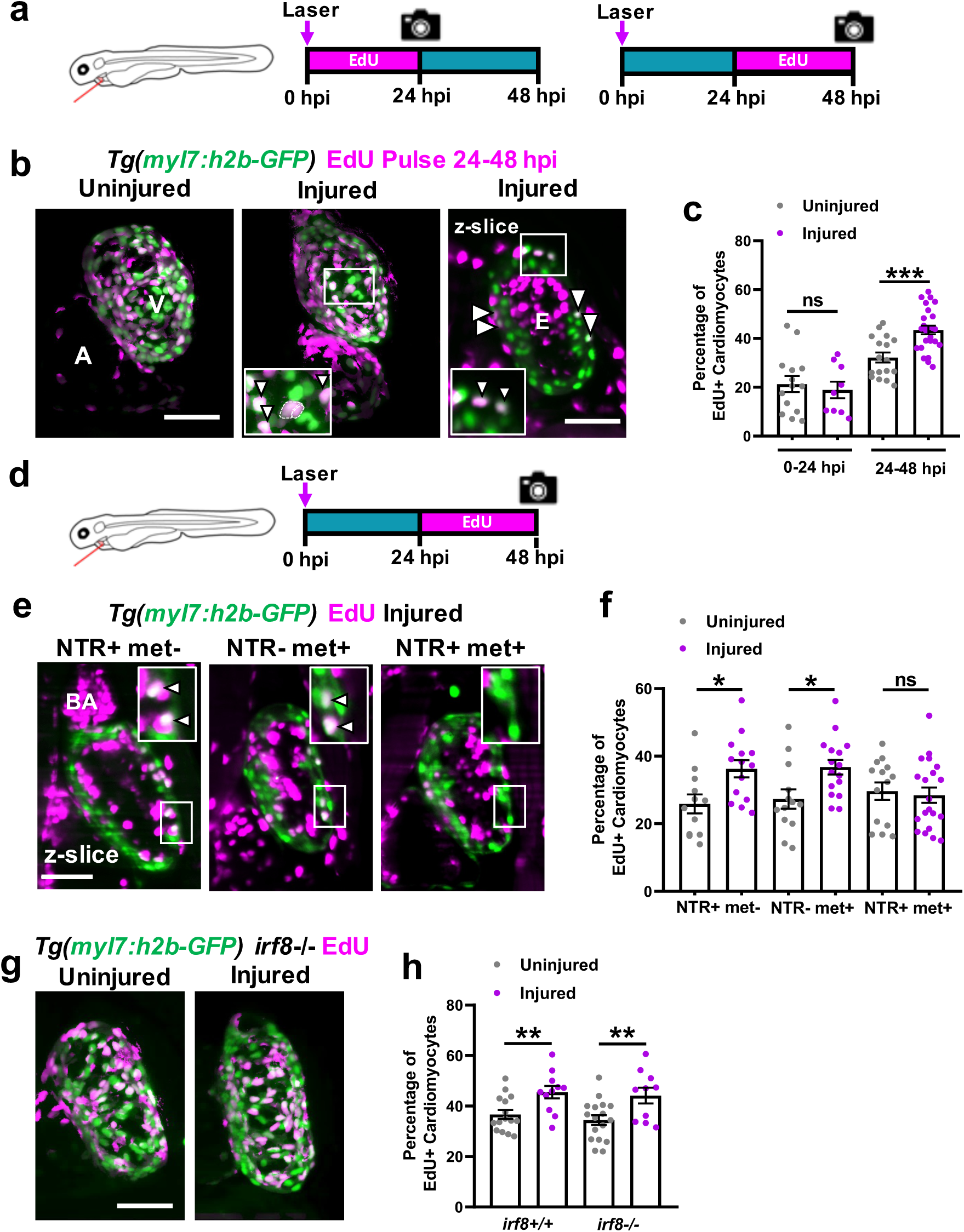
Macrophage ablation abolishes injury-dependent cardiomyocyte proliferation. (a) Schematic illustrating EdU pulse strategy for labelling proliferating cardiomyocytes over 0-24 hpi (left) and 24-48 hpi (right). (b) Representative images of EdU-stained hearts from *Tg(myl7:h2b-GFP)* at 48 hpi. Non-myocardial EdU signal is excluded post-acquisition to allow interpretable maximal intensity projections (MIPs). A = atrium, v = ventricle; white boxes = zoom panels; white arrowheads = EdU+ cardiomyocyte nuclei and dashed line = outline of dividing cardiomyocyte daughter nuclei. (c) Quantification of the percentage of ventricular EdU+ cardiomyocytes in uninjured and injured *Tg(myl7:h2b-GFP)* hearts pulsed over 0-24 hpi or 24-48 hpi. \*\*\**p≤0.001* unpaired t test. (d) Schematic illustrating EdU pulse strategy for labelling proliferating cardiomyocytes over 24-48 hpi in *Tg(myl7:h2b-GFP;csfr1a:NfsB-mCherry)* larvae per standard macrophage ablation groups. (e) Representative images of EdU-stained hearts from *Tg(myl7:h2b-GFP;csfr1a:NfsB-mCherry)* acquired by light-sheet microscopy at 48 hpi. White boxes = zoom panels; white arrowheads = EdU+ cardiomyocyte nuclei and BA = bulbous arteriosus. (f) Quantification of the percentage of ventricular EdU+ cardiomyocytes in uninjured and injured *Tg(myl7:h2b-GFP;csfr1a:NfsB-mCherry)* hearts pulsed over 24-48 hpi. \**p≤0.05* Kruskal-Wallis test and Dunn’s multiple comparison post-hoc test. (g) Representative images of uninjured and injured EdU-stained hearts from *irf8^-/-^ Tg(myl7:h2b-GFP)* acquired by light-sheet microscopy at 48 hpi. Non-myocardial EdU signal is excluded post-acquisition to allow interpretable maximal intensity projections. (h) Quantification of the percentage of ventricular EdU+ cardiomyocytes in uninjured and injured *irf8^+/+^* and *irf8^-/-^ Tg(myl7:h2b-GFP)* hearts pulsed 24-48 hpi. All images are maximum intensity projections of 3D LSFM stacks, unless otherwise stated. Scale bars = 50 μm. \*\**p≤0.01* unpaired t test

To understand if macrophages are required for the injury-dependent increase in cardiomyocyte proliferation, EdU was pulsed during the proliferative 24-48 hpi window in the macrophage-less models (Figure 4d). In the metronidazole-nitroreductase ablation model we found that the percentage of EdU+ cardiomyocytes increased in injured hearts in both the NTR+met- and NTR-met+ control groups, but not in the macrophage-ablated NTR+met+ group (Figure 4e & 4f). This result indicates that macrophages are a requirement for injury-dependent increase in cardiomyocyte proliferation. However, in contrast to the metronidazole-nitroreductase ablation model, analysis of cardiomyocyte proliferation in *irf8^-/-^* mutants revealed that they too significantly increased the percentage of EdU+ cardiomyocytes following injury, comparably to *irf8^+/+^* larvae (Figure 4g & 4h).

To resolve this disparity, we examined more closely the differences between these models. We found, like others^18^, that *irf8^-/-^* mutants possess a greater global number of neutrophils than observed in *irf8^+/+^* fish and mount a larger neutrophil response to injury (Supplementary Figure 5b & 5c). Since we do not observe an increased neutrophil response in NTR+met+ larvae, we hypothesised that neutrophils might be compensating for macrophages in *irf8^-/-^* larvae (Supplementary Figure 5d). To test this hypothesis, we inhibited neutrophil recruitment in *irf8^-/-^* larvae using the receptor antagonist ‘SB225002’ which blocks CXCR1/2 activation, a key chemokine receptor for neutrophil migration. CXCR1/2 inhibition successfully lowered the number of recruited neutrophils (2.0±3.4 vs 0.43±0.18) and abolished the injury-associated increase in cardiomyocytes in *irf8^-/-^* (Supplementary Figure 5e-g). Taken together, this suggests that macrophages are required for cardiomyocyte proliferation but can be substituted by excess neutrophils.

### Regenerating larval hearts resolve inflammation and enter a reparative stage by 48 hpi

Next, we sought to understand which biological processes might still be occurring by the final 48 hpi timepoint of the larval cardiac injury model. We performed RNAseq on pooled, uninjured and injured larval hearts at 48 hpi (Figure 5a). We found 418 genes were upregulated (log_2_ fold change >1), and 1,046 downregulated in injured hearts. We did not observe differential expression of markers of proliferation such as MCM2, mKi67 and PCNA, suggesting that the proliferation we observe from 24 hpi is concluded by 48 hpi (Figure 5b). In agreement with this, gene ontology analysis indicated categories such as growth factors and cell proliferation not to be enriched at 48 hpi (Supplementary Figure 6e).

**Figure 5:**
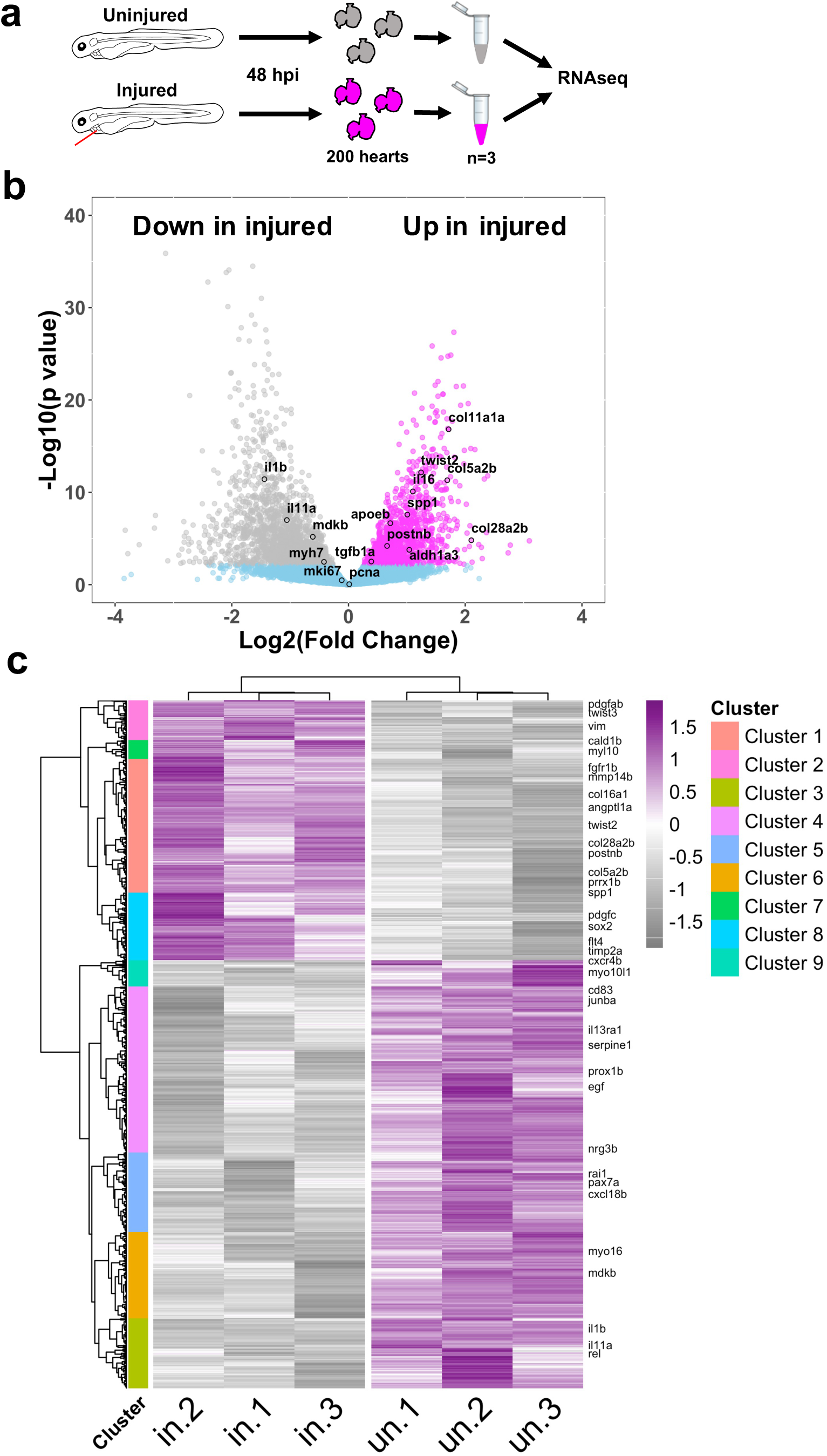
Bulk RNAseq analysis of larval hearts following injury. (a) Schematic illustrating the extraction of uninjured and injured hearts at 48 hpi and the pooling of 200 hearts per biological replicate for RNAseq, n=3. (b) Volcano plot showing the Log_2_(Fold Change) and –Log_10_(p value) for transcripts of each detected gene. Genes whose adjusted p values fall below 0.05 are deemed statistically non-significant and coloured blue. Genes up regulated in injured hearts are coloured magenta and those upregulated in uninjured hearts are coloured grey. (c) Heatmap displaying statistically significantly differentially expressed genes with a Log_2_(Fold Change) >0.5. Genes were hierarchically clustered by Pearson correlation with z scaling. Clusters are indicated on the left with their dendrogram. Magenta = high expression; grey = low expression. Genes with relevance to cardiac regeneration are highlighted as annotations on the right of the plot. n=3

Most inflammatory and M1 markers were either not differentially expressed or were downregulated in injured hearts, such as Il1b (Figure 5b & Supplementary file 1). In contrast, we found injury-associated upregulation of 39 collagen isoforms, several profibrotic genes such as *tgfb1a* and markers of epithelial to mesenchymal transition (EMT) such as *vimentin*. Similarly, hierarchical clustering of differentially expressed genes revealed 9 distinct clusters with Cluster 1 being upregulated in injured hearts and enriched in collagens, matrix metalloproteins (MMPs) and fibroblast growth factors (FGFs) (Figure 5c, Supplementary file 2). Additionally, Cluster 2 contained several EMT genes, Cluster 8 genes relating to cell recruitment and lymphangiogeneis whilst Cluster 7 contained several embryonic-associated myosins and myosin binding proteins such as *myl10* and *cald1b*. Clusters 3-6 & 9 were downregulated in injury, Cluster 2 was enriched for immune genes and Clusters 4 and 6 for growth factors, with Clusters 3 and 5 having no clear identity. Taken together, our RNAseq results suggest that the inflammatory and proliferative stages are largely concluded by 48 hpi and that a pro-resolving and reparative phase dominates thereafter.

### Cardiac injury induces epicardial activation and VEGFaa upregulation

Our detailed characterisation of the larval laser injury model revealed a macrophage-dependent, cardiomyocyte proliferative response occurring at 24-48 hpi. We therefore utilised the rapidity and imaging opportunities offered by the model to investigate the underlying mechanism of the induction of cardiomyocyte proliferation. Epicardial VEGFaa has recently been demonstrated to drive cell cardiomyocyte proliferation in adult zebrafish following cryoinjury and we hypothesised the same mechanism might drive cardiomyocyte proliferation in the injured larval heart^33^.

We found robust *vegfaa:GFP* expression specifically in mesothelial cells overlying the myocardium (Figure 6a). Colocalisation with established epicardial marker *tcf21* in uninjured *Tg(tcf21:DsRed;vegfaa:GFP)* larvae confirmed these cells to be early epicardium (Figure 6b). Next we investigated if epicardial *vegfaa:GFP* expression changes following injury by 3D fluorescence intensity analysis of uninjured and regenerating hearts. We found that epicardial *vegfaa:GFP* intensity increased significantly at 48 hpi (Figure 6c & 6d). Interestingly, this was due both to an increase in the number of epicardial cells and their individual intensity suggesting the epicardium activates and responds to injury by both proliferation and gene expression changes to increase VEGFaa (Supplementary figure 7c & 7d).

**Figure 6:**
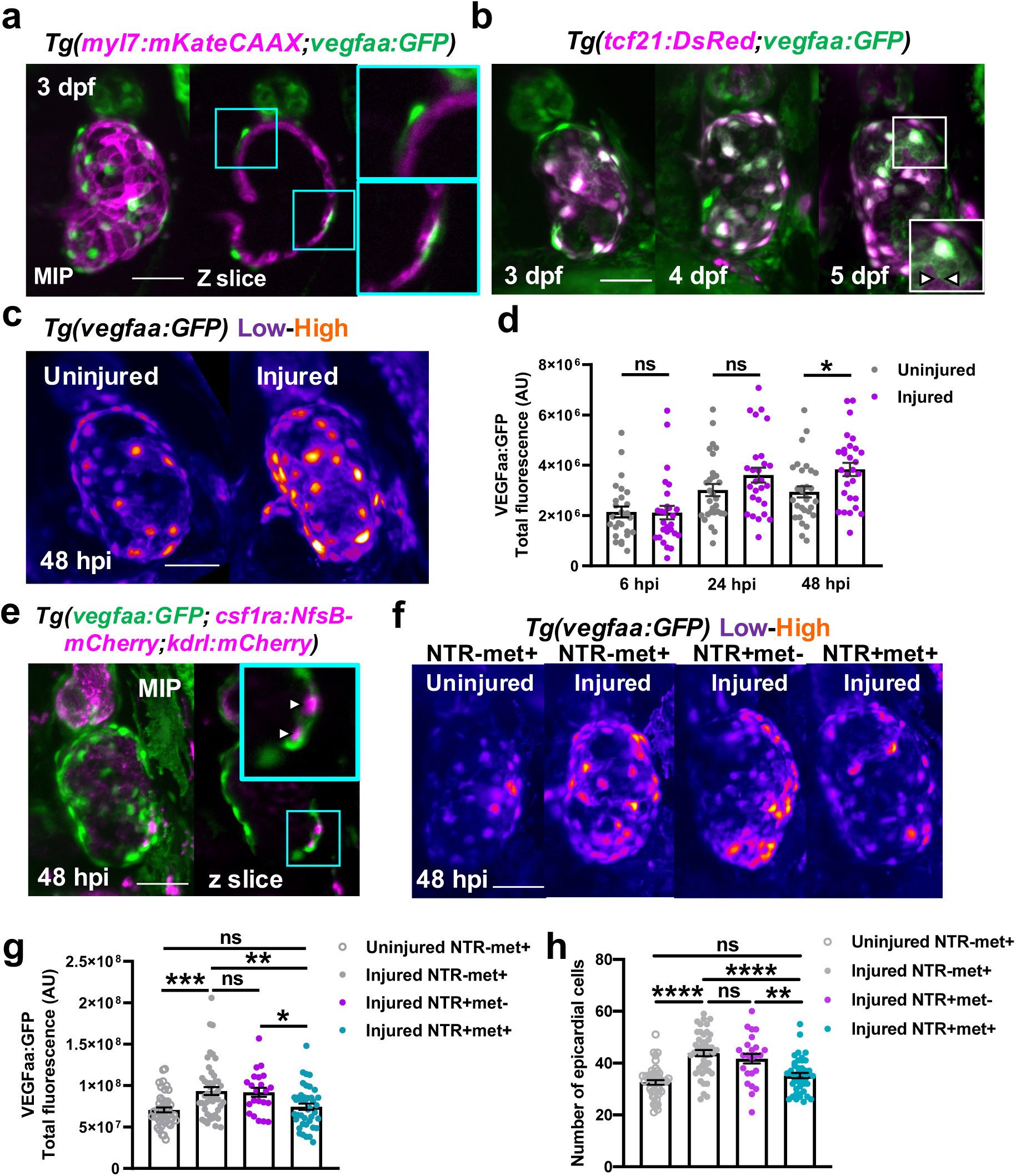
Macrophages stimulate epicardial cell number expansion following cardiac injury. (a) Representative LSFM image of an uninjured 3 dpf ventricle from a *Tg(myl7:mKateCAAX;myl7:h2b-GFP)* larva showing vegfaa+ cells (green) overlying myocardium (magenta). Cyan box = zoom panel. (b) Representative images of 3, 4 and 5 dpf ventricles from a *Tg(tcf21:DsRed;vegfaa:GFP)* larvae acquired by LSFM, showing high colocalization of vegfaa with epicardial marker tcf21. White arrowheads = heterogenous marker expression and white box = zoom panel. (c) Representative images of uninjured and injured ventricles from *Tg(vegfaa:GFP)* larvae acquired at 48 hpi by LSFM. “Heat” LUT applied to highlight increased intensity of epicardial vegfaa:GFP in injured hearts. (d) Quantification of total ventricular VEGFaa:GFP fluorescence in uninjured and injured hearts over standard injury model timepoints, n=28-30. \**p≤0.05* One way ANOVA followed by Holms-Sidak’s multiple comparison Post-hoc tests. (e) Representative image of a ventricle from a *Tg(vegfaa:GFP;csfr1a:NfsB-mCherry;kdrl:hsa.HRAS-mCherry)* (abbreviated to kdrl:mCherry) larva at 48 hpi showing macrophages in the epicardial-myocardial niche (white arrowheads). Cyan box = zoom panel. (f) Representative LSFM images of uninjured and injured ventricles from *Tg(vegfaa:GFP;csfr1a:NfsB-mCherry)* larvae from metronidazole-nitroreductase macrophage ablation groups at 48 hpi. “Heat” LUT is applied to highlight increase in overall fluorescence in injured groups except NTR+met+. (g) Quantification of total vegfaa:GFP fluorescence (g) and epicardial cell number (h) in uninjured and injured ventricles from *Tg(vegfaa:GFP;csfr1a:NfsB-mCherry)* larvae from metronidazole-nitroreductase macrophage ablation groups at 48 hpi. All images are maximum intensity projections of 3D LSFM stacks. Scale bars = 50 μm, n=46. \**p≤0.05,* \*\**p≤0.01,* \*\*\**p≤0.001* One way ANOVA followed by Holms-Sidak’s multiple comparison Post-hoc tests.

### Macrophages localise to the epicardial niche and induce the expansion of epicardial cell number

Given that our data showed that macrophage ablation abolishes injury-dependent cardiomyocyte proliferation (Figure 5f), we hypothesised that macrophages might be required for epicardial activation. To test this hypothesis, we ablated macrophages and assessed if epicardial activation still occurred at 48 hpi. Following injury we observed increased *vegfaa:GFP* expression in both macrophage-replete NTR-met+ and NTR+met- groups, but not in macrophage ablated NTR+met+ hearts (Figure 6f & 6g). Interestingly, macrophage ablation did not affect *vegfaa:GFP* expression per cell, but did block the expansion of epicardial cell number following injury (Figure 6h & Supplementary Figure 7e). Furthermore, 3D analysis of macrophage localisation following injury showed that recruited macrophages invade the myocardial-epicardial niche and synapse with epicardial cells (Figure 6e). Importantly, macrophage or neutrophil *vegfaa:GFP* expression was not observed at any timepoint (Supplementary Figure 7a & 7b). Our data therefore strongly suggest that the recruitment of macrophages to the epicardium is essential for subsequent epicardial activation, thus increasing net cardiac *vegfaa* expression.

### VEGFaa is both required and sufficient for cardiomyocyte proliferation in larval zebrafish

To verify if epicardial VEGFaa was driving cardiomyocyte proliferation in larval cardiac regeneration, we first tested if VEGFaa was sufficient to stimulate cardiomyocyte proliferation. Recombinant zebrafish VEGFaa protein (zfVEGFaa) was intravenously microinjected into the circulation of 72 hpf *Tg(myl7:h2b-GFP)* larvae and total cardiomyocyte number assessed at 24 and 48 hpt (hours post-treatment) (Figure 7a). zfVEGFaa increased total cardiomyocyte number by 13.3% relative to PBS-injected controls at 24 hpt (Figure 7b & 7c).

**Figure 7:**
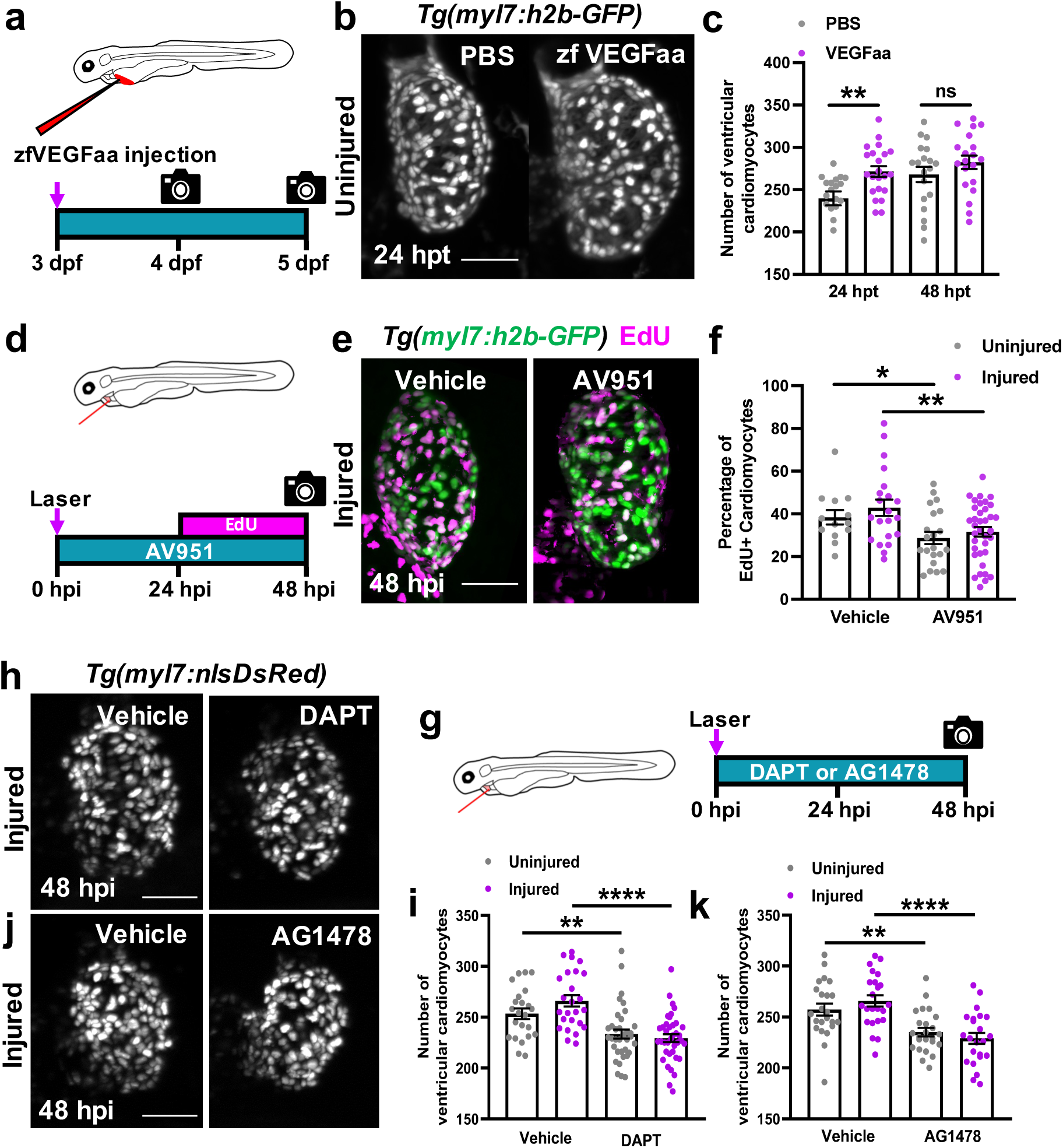
VEGFaa signalling is both sufficient and necessary to drive cardiomyocyte proliferation. (a) Schematic illustrating zfVEGFaa treatment strategy via microinjection into the common cardinal vein of uninjured larvae at 72 hpi. (b) Representative LSFM images of *Tg(myl7:h2b-GFP)* larvae at 24 hpi treated with PBS 0.1% BSA or zfVEGFaa 0.1% BSA injection. (c) Quantification of ventricular cardiomyocyte number in *Tg(myl7:h2b-GFP)* larvae at 24 and 48 hpi treated with PBS 0.1% BSA or zfVEGFaa 0.1% BSA injection, n=20. \*\**p≤0.01* unpaired t test. (d) Schematic illustrating AV951 treatment and EdU pulsing strategy for uninjured and injured larvae. (e) Representative images of injured ventricles from *Tg(myl7:h2b-GFP)* larvae, EdU stained and bathed in vehicle or AV951, imaged at 48 hpi by LSFM. Non-myocardial EdU signal is excluded post-acquisition to allow interpretable maximal intensity projections (MIPs). (f) Quantification of the percentage of EdU+ cardiomyocyte nuclei from uninjured and injured ventricles from *Tg(myl7:h2b-GFP)* larvae, EdU stained and bathed in vehicle or AV951, n=13-36. \**p≤0.05,* \*\**p≤0.01* unpaired t test. (g) Schematic illustrating the treatment strategy for DAPT and AV951 bathing of uninjured and injured larvae. (h) Representative images of injured *Tg(myl7:nlsDsRed)* larvae treated with vehicle or DAPT, acquired at 48 hpi by LSFM. (i) Quantification of ventricular cardiomyocyte number in uninjured and injured *Tg(myl7:h2b-GFP)* larvae at 48 hpi treated with vehicle or DAPT, n=24-40. \*\**p≤0.01,* \*\*\*\**p≤0.0001* Unpaired t test.(j) Representative images of injured *Tg(myl7:nlsDsRed)* larvae treated with vehicle or AG1478, acquired at 48 hpi by LSFM. (k) Quantification of ventricular cardiomyocyte number in uninjured and injured *Tg(myl7:h2b-GFP)* larvae at 48 hpi treated with vehicle or AG1478, n=24. All images are maximum intensity projections of 3D LSFM stacks. \*\**p≤0.01,* \*\*\*\**p≤0.0001* Unpaired t test. Scale bars = 50 μm.

To test if VEGF signalling is required for injury-associated cardiomyocyte proliferation, we used a high-affinity, pan-VEGFR receptor antagonist AV951 (Tivozanib) to block VEGF signalling^34^. We bathed larvae in 10nM AV951 over the course of our cardiac injury model, pulsed with EdU at 24-48 hpi and quantified EdU+ cardiomyocytes at 48 hpi (Figure 7d). Interestingly, AV951 decreased the proportion of EdU+ cardiomyocytes in both the uninjured and injured groups (uninjured 38.4±3.4 vs 28.0±3.0 and injured 42.9±3.8 vs 31.6±2.2, n=13-36) (Figure 7e & 7f). Together, these data suggest that VEGF signalling in the heart is driving cardiomyocyte proliferation in the larval heart, both as part of normal development and following cardiac injury.

### Notch and Nrg-ErbB signalling are required for cardiomyocyte proliferation

We next investigated if macrophage-induced epicardial VEGFaa signalling could be interacting with more established effectors of cardiomyocyte proliferation. Notch and Nrg-ErbB were strong candidates as both are required for adult heart regeneration and cardiomyocyte proliferation in adult zebrafish^35–38^. We first verified if these signalling pathways were required for cardiomyocyte proliferation in the larval heart following injury. We laser-injured *Tg(myl7:nlsDsRed)* larvae, and bathed them in 100μM of pan-notch inhibitor DAPT ((N-[N-(3,5-difluorophenacetyl)-l-alanyl]-S-phenylglycine t-butyl ester) (Figure 7g). DAPT is a gamma secretase inhibitor and has been demonstrated in both zebrafish and drosophila to phenocopy notch mutants^39–41^. Notch signalling inhibition decreased cardiomyocyte number by ∼8% in uninjured hearts (253.3±5.2 vs 233.4±4.5, n=25-38) and ∼14% injured hearts (266.0±5.6 vs 229.5±3.6, n=25-38). (Figure 7h & 7i).

We repeated this experiment, with 1.75μM ErbB2 antagonist AG1478 (Figure 7g). Small molecule inhibitor AG1478 selectively inhibits ErbB2, a required co-receptor for ErbB4 dimerization and subsequent neuregulin signal transduction and faithfully phenocopies *erbb2* mutants^42^. Interestingly, the results exactly replicated those of the notch signalling inhibition experiment, decreasing cardiomyocyte number by ∼9% in uninjured (257.2±5.8 vs 235.2±4.2) and by ∼13% in injured hearts (265.7±5.5 vs 229.5±5.9) (Figure 7j & 7k). These results confirm that both notch signalling and Nrg-ErbB signalling are required for the expansion of cardiomyocyte number in both uninjured and injured larval hearts.

### Cardiac injury and VEGFaa induce endocardial notch signalling

Given the individual requirement of VEGF, notch and Nrg-ErbB signalling for cardiomyocyte proliferation in the larval heart, we sought to understand if these signalling components might act in one pathway. Previous studies have demonstrated developmental larval zebrafish heart growth to be activated by cardiac contraction, via endocardial-notch>Nrg-ErbB signalling^43, 44^. We hypothesised that VEGFaa might be driving cardiomyocyte proliferation by increasing endocardial notch signalling and consequently augmenting this developmental pathway (Figure 8a).

**Figure 8:**
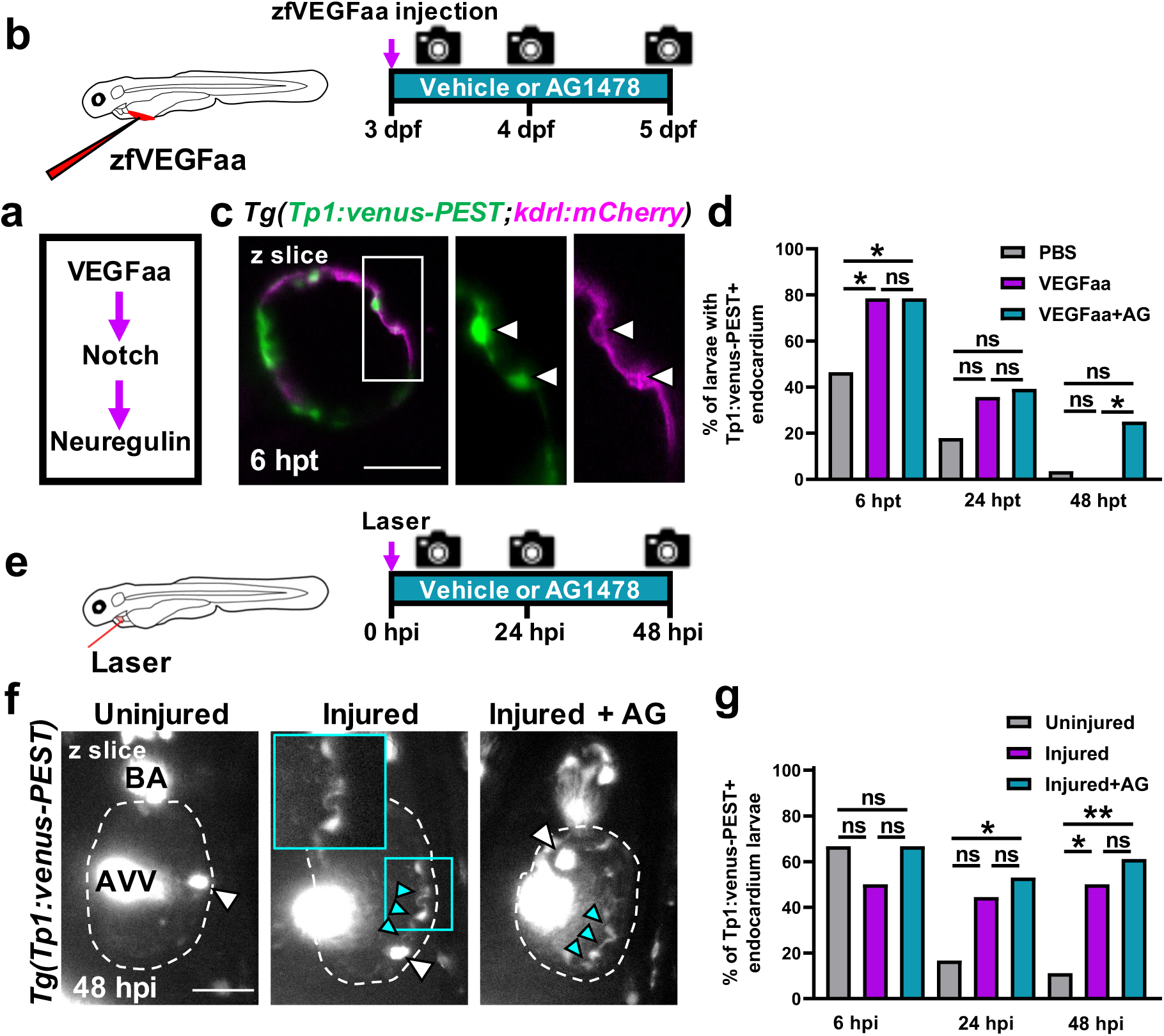
VEGFaa drives cardiomyocyte proliferation by endocardial notch signalling. (a) Hypothesised signalling pathway active in uninjured and injured larval hearts driving cardiomyocyte proliferation. (b) Schematic illustrating the treatment strategy for the injection of uninjured larvae with zfVEGFaa and continuous bathing in AG1478 solution. (c) Representative LSFM-acquired z plane showing notch expression colocalising with endocardium in *Tg(Tp1:venus-PEST;kdrl:hsa.HRAS-mCherry),* abbreviated in the figure to *Tg(Tp1:venus-PEST;kdrl:mCherry).* AG1478 abbreviated to AG, white box = zoom panel. (d) Quantification of the proportion of larvae with notch+ endocardium at 6, 24, and 48 hpt following zfVEGFaa injection and bathing in AG1478, n=28. \**p≤0.05* Fisher’s exact test. (e) Schematic illustrating the treatment strategy for the lasering of larvae and continuous bathing in AG1478 solution. (f) Representative z plane images of uninjured, injured and injured AG-treated ventricles from *Tg(tp1:venus-PEST)* larvae acquired by light-sheet microscopy at 48 hpi. BA = bulbous arteriosus; AVV = atrioventricular valve; white arrowheads =laterally inhibited cardiomyocytes, cyan arrowheads = notch+ endocardium; cyan box = zoom panel. (g) Quantification of the proportion of larvae with notch+ endocardium at 6, 24, and 48 hpt following laser injury and bathing in AG1478, n=18. All images are maximum intensity projections of 3D LSFM stacks unless otherwise stated. \**p≤0.05,* \*\**p≤0.01* Fisher’s exact test. Scale bars = 50 μm.

To test if VEGFaa could activate endocardial notch signalling, we utilised the notch signalling reporter line *Tg(Tp1:venus-PEST)* as a readout of cardiac notch signalling. Recombinant zfVEGFaa injected into 3 dpf larvae, and their hearts were analysed *via* heart-synchronised light-sheet microscopy at 6, 24 and 48 hpt (Figure 8b). Furthermore, an additional group of larvae were injected with zfVEGFaa but also bathed in ErbB2 antagonist AG1478. According to our hypothesised pathway (Figure 8a), we reasoned that zfVEGFaa injection should upregulate notch signalling but that inhibition of Nrg-ErbB signalling should be unable to suppress zfVEGFaa-induced notch upregulation.

Notch signalling was primarily in the endocardium, colocalising with endothelial reporter *kdrl:mCherry* but was relatively low intensity and only detectable in a subset of larvae at any given timepoint (Figure 8c). We found, zfVEGFaa injection increased the percentage of larvae with notch+ (Tp1:Venus+) endocardium (46.4% to 78.6%, n=28) at 6 hpi but not at the 24 and 48 hpi timepoints (Figure 8d). Furthermore, AG1478 failed to block the increase in the percentage of notch+ hearts following zfVEGFaa injection, confirming Nrg-ErbB signalling was not upstream of *vegfaa* or notch signalling. In fact, zfVEGFaa+AG1478 treated larvae had a substantially higher percentage of notch+ hearts at 48 hpi than those treated with zfVEGFaa alone (25.0% vs 0%). This is suggestive of a negative feedback mechanism, supporting previous findings of Nrg-ErbB being downstream of notch signalling^43^.

To test if this pathway (Figure 8a) acted similarly in injury, we substituted zfVEGFaa injection for cardiac injury and repeated the experiment (Figure 8e). Cardiac injury similarly increased the percentage of hearts possessing notch+ endocardium but this occurred later, at 48 hpi (50.0% vs 11.1%, n=18) (Figure 8f & 8g). As before, AG1478 did not block notch signalling, rather it seemed to enhance it. Whilst the percentage of notch+ hearts in the injured group did not significantly increase by 24 hpi, injured+AG1478 treated larvae did significantly increase relative to uninjured larvae (52.9% vs 11.1%).

Taken together, these results demonstrate that cardiac injury and VEGFaa increase endocardial notch signalling, providing a novel mechanism whereby macrophages can trigger cardiomyocyte proliferation via stimulation of epicardial *vegfaa* expression.

## Discussion

In this study we have presented the first detailed characterisation of the larval zebrafish model of heart regeneration, demonstrating the heterogeneity and plasticity of macrophages in cardiac injury and testing the requirement of macrophages for the removal of apoptotic cells, cardiomyocyte proliferation, epicardial activation and recovery of cardiac structure and function. Furthermore, we demonstrated the utility of the larval cardiac injury model by taking advantage of its *in vivo* cardiac imaging opportunities and amenability to pharmacological intervention to discover a novel role for macrophages in driving cardiomyocyte proliferation via epicardial activation.

Our examination of macrophages in larval zebrafish cardiac injury suggests that they may faithfully recapitulate the phenotypic complexity and function found in the adult cryoinjury model. As previously shown in adult hearts^29^, we detected mpeg1+csf1ra+ and mpeg1+csf1ra- macrophage subsets. We found these cells to have identical recruitment dynamics and no obvious differences in morphology or behaviour. Recent Cre-Lox lineage tracing has shown that mpeg1+csf1ra- cells have a non-haematopoietic origin, are csfr1a-independent developmentally and, unlike mpeg1+csf1ra+ cells, are not phagocytic^45, 46^. Future work should focus on understanding the precise roles of these subsets in cardiac regeneration, in particular mpeg1+csf1ra- macrophages.

In addition to macrophage heterogeneity, we observed macrophage phenotypic plasticity. We used heartbeat-synchronised live imaging to show that macrophages can convert from mpeg1+tnfa-to mpeg1+tnfa+. This is the first time that macrophage phenotype conversion has been imaged in the heart. Studies examining zebrafish macrophages in spinal cord and tail transection have demonstrated *tnfa* to mark M1- like macrophages, which then transition to M2-like macrophages^24^. Our success in increasing the percentage of tnfa+ macrophages by canonical M1-polarising cytokine IFN-γ-rel suggests that early tnfa+ macrophages are indeed proinflammatory. In agreement with findings in the in the adult cryoinjured heart^29^, we found this tnfa+ population of macrophages to be transient, only observed in the early response at 24 hpi. Similarly, our RNAseq data showed that, by 48 hpi, injured larval hearts downregulate inflammatory cytokines and growth factors but upregulate collagens and reparative cytokines. Our finding that a pro-resolving, fibrotic program is activated in injured hearts, despite full structural and functional recovery, is in agreement with a recent study showing the scar-deficient runx1^-/-^ zebrafish to undergo successful cardiac regeneration^47^. It might be that the fibrotic program is concomitantly activated upon resolution of inflammation, irrespective of a requirement for scar tissue.

We used two separate methods to examine the role of macrophages in larval heart regeneration. Interestingly, ablation via csf1ra-driven nitroreductase expression was still able to abolish numbers of csf1ra-macrophages at the injured heart. Possibly this is indicative of a positive-feedback system where csf1ra-macrophage recruitment is dependent on csf1ra+ macrophages. Cell death data acquired by either technique demonstrated that macrophages are required for the timely removal of apoptotic cells following injury. Interestingly, these apoptotic cells do eventually seem to be cleared even in the absence of macrophages. Our live imaging showed that dead cardiomyocytes can be expelled from the myocardium independently of macrophages. It is possible that this is a mechanical consequence of cardiac contraction, although a similar phenomenon is known to occur in neuroepithelium where neurons appear to extrude apoptotic cells out of tissue^48^.

Surprisingly, we found that the absence of macrophages did not affect the structural or functional recovery of the injured larval heart, despite macrophages being required for cardiomyocyte proliferation. This is in contrast to past studies where liposomal clodronate macrophage ablation and CCR2-antagonist inhibition of macrophage recruitment in regenerative neonatal mice, adult zebrafish and axolotl hearts causes blocked or delayed resolution of the infarct area^15, 16, 49^. The contrasting results in the larval heart might simply be a consequence of its small size and low transmural pressure, allowing surviving myocardium to rapidly ‘knit’ back together. Supporting this, we observed individual cardiomyocytes extending protrusions into the lesion. Previous histological analysis of the border zone in injured adult zebrafish and neonatal mouse hearts has shown cardiomyocytes exhibiting a similar mesenchymal phenotype following partial dedifferentiation and disassembly of sarcomeres^50, 51^. However, this is the first time this behaviour has been verified by time-lapse imaging live in a beating heart.

Studies in adult zebrafish and neonatal mice have shown ablation of macrophages to decrease cardiomyocyte proliferation^15, 17^. However, early revascularisation is critical for cardiomyocyte proliferation and is macrophage-dependent, calling into question whether macrophages directly induce cardiomyocyte proliferation^17, 52^. Larval hearts do not have supporting vasculature^26^; therefore our finding that cardiomyocyte proliferation is macrophage-dependent suggests macrophages facilitate cardiomyocyte proliferation by means other than revascularisation. Indeed, our data indicate a novel mechanism whereby macrophages are recruited to the epicardial-myocardial niche and induce expansion of epicardial cell numbers and increase in the expression of mitogenic VEGFaa. This might explain previous findings in developing and injured mouse hearts where yolk-derived and Gata6+ pericardial cavity macrophages are recruited to the epicardium, respectively^53, 54^ Future studies should seek to identify precisely how macrophages activate epicardium. Given we found that neutrophils can compensate for macrophages for cardiomyocyte proliferation, it is possible that a shared inflammatory factor triggers epicardial activation.

Endocardial notch signalling is required for cardiomyocyte proliferation in cryoinjured adults and myocardial growth by downstream Nrg1 in larvae^43, 55, 56^. Therefore, our finding that injury and VEGFaa increase endocardial notch signalling reveals an important mechanism whereby the epicardium can induce cardiomyocyte proliferation. In agreement with previous studies, we showed both notch and Nrg-ErbB signalling to be required for expansion of cardiomyocyte numbers in both uninjured and injured hearts^57–59^. Since we found VEGFaa inhibition decreased cardiomyocyte proliferation in uninjured larval hearts, it is likely that VEGFaa>notch>Nrg-ErbB is a developmental cardiac growth pathway that is upregulated upon injury and thus might be conserved in mammals. Our discovery that macrophages act upstream of this pathway therefore opens up exciting immunomodulatory opportunities for therapeutic enhancement of cardiac repair in the future.

## Materials and Methods

### Zebrafish husbandry and lines used

Zebrafish husbandry and maintenance was conducted as per standard operating procedures, in accordance with the Animals (Scientific Procedures) Act, 1986 and approved by The University of Edinburgh Animal Welfare and Ethical Review Board in a United Kingdom Home Office-approved establishment. All our experiments were performed on staged zebrafish aged between 3 dpf and 5 dpf. The following transgenic and mutant lines were used: *Tg(myl7:eGFP)^twu^*^26 60^, *Tg(mpx:mCherry)^uwm^*^7 61^, *Tg(mpeg1:mCherry)^gl^*^23 62^, *Tg(mpeg1:eGFP)^gl22^* ^62^, (*Tg(mpx:GFP) ^i114^* ^63^, *Tg(myl7:h2b-GFP)^zf52^* ^64^, *Tg(myl7:mKateCAAX)^SD11^* ^65^, *Tg(fms:Gal4.VP16)^i186^,* referred to as csfr1a:gal4^66^, *Tg(UAS-E1b:NfsB-mCherry)^c264^* abbreviated to UAS:NfsB-mCherry^67^*, Tg(vegfaa:eGFP)^PD260^* ^33^*, Tg(myl7:nlsDsRed)^f^*^2 68^ *Tg(TNFa:eGFP)^sa43296^* ^24^, *Tg(Tp1:venus-PEST)^S940^* ^69^*, Tg(kdrl:hsa.HRAS-mCherry)^S896^* ^70^, *Tg(kdrl:GFP)^la116^* ^71^*, Tg(tcf21:DsRed)^PD37^* ^72^, *Tg(myl7:gal4:myl7:GFP^)cbg2Tg^* ^73^ and irf8^st95/st95 27^ referred to as *irf8^-/-^. Tg(csf1ra:gal4:UAS:NfsB-mCherry)* is abbreviated to csf1ra:NfsB-mCherry throughout the manuscript for simplicity. Adults were day-crossed as appropriate to yield desired combinations of transgenes in embryos. Embryos were treated with 0.003% phenylthiourea (Fisher Scientific) at 7 hpf to prevent pigment formation and therefore enhance image clarity. Embryos and larvae were incubated at 28.5°C in conditioned media/water (6.4 mM KCl, 0.22 mM NaCl, 0.33 mM CaCl_2_·2H_2_O, 0.33 mM MgSO4·7H_2_O) + 0.1% methylene blue (w/v) and imaged at room temperature (23°C) using epifluorescence or light sheet fluorescence microscopy (details below). When necessary, larvae were anesthetized using 40 μg/ml tricaine methanesulfonate (Sigma Aldrich) in conditioned media.

### Cardiac laser injury

A Zeiss Photo Activated Laser Microdissection (PALM) laser system (Zeiss) was used to precisely cause a localised injury at the ventricular apex of anesthetized 72 hpf larvae^26^. Larvae were mounted on a glass slide in 20 μl anesthetized conditioned media and lasered via a 20X objective. Injuries were deemed successful and complete once ventricular contractility decreased, the apex had shrunk, and the myocardial wall had swollen without causing cardiac rupture and subsequent bleeding. A successful cardiac injury results in the portion of dysfunctional tissue losing fluorescent myocardial transgenic fluorescence signal. Uninjured larvae were treated in the same manner up to the point of laser injury, when they were individually transferred into single wells of a 24-well plate and maintained in the same environmental conditions as injured fish.

### Epifluorescence microscopy

Larvae were mounted laterally in conditioned media on a glass slide and imaged using a Leica M205 FA stereomicroscope with GFP and mCherry filters. For all serial timepoint epifluorescence imaging experiments, number of immune cells on the heart were quantified by manually observing and counting cells moving synchronously with the beating heart. Heart images were acquired using 2X 0.35NA objective.

### Heart-synchronised light-sheet microscopy

Individual larvae were prepared for light sheet fluorescence microscopy (LSFM) by embedding in 1% low melting-point agarose (ThermoFisher) in anesthetized conditioned media inside FEP tubes (Adtech Polymer Engineering). Agar embedding prevents gradual drift of the embryo in the FEP tube, without causing developmental perturbations during long-term imaging. Larvae were used only once for a time-lapse imaging experiment, and any repeats shown come from distinct individuals. Larvae were mounted head down such that the heart faces toward both illumination and imaging objectives to improve image clarity. All LSFM experiments were performed at room temperature (23°C). Camera exposure times ranged from 5-15 ms, laser excitation power was 11mW and scans were performed at 3-5 minute intervals. Brightfield images acquired at 80 fps were analysed in real-time to enable optically-gated acquisition of fluorescence z slices at a set phase of cardiac contraction, usually mid diastole. The setup of our custom-built LSFM system has been previously reported in detail^30^.

### Metronidazole-nitroreductase macrophage ablation model

In order to selectively ablate macrophages prior to cardiac injury, embryos were incubated as previously described until 48 hpf and then treated as follows. Embryos were carefully dechorionated at 48 hpf and screened based on fluorescence and split into groups appropriate to the experiment, for example larvae were always split into csf1ra:gal4;UAS:NfsB-mCherry+ and csf1ra:gal4;UAS:NfsB-mCherry-. Embryos were then transferred to either conditioned water or a 0.5mM metronidazole (Thermo Fisher Scientific) solution, both solutions also contained 0.003% phenylthiourea (Thermo Fisher Scientific) and 0.2% DMSO (Sigma Aldrich). Larvae were then incubated in these solutions in the dark at 28. 5°C for 24 hours prior to injury at 72 hpf. Larvae were then removed from the metronidazole solution and vehicle solution and placed in fresh conditioned water + 0.003% phenylthiourea for the remainder of the experiment. As shown in Figure 2, this is sufficient to ablate macrophages prior to injury and completely block subsequent macrophage recruitment to the injured heart.

### Neutral red staining

Larvae were incubated at 72 hpf in 5μg/mL neutral red in conditioned water for 5 hours in the dark at 28.5°C. Larvae were then washed twice for 5 minutes in conditioned water, anaesthetised with 40 μg/ml tricaine methanesulfonate and imaged by brightfield microscopy on a Leica M205 FA stereomicroscope.

### Genotyping of *irf8^-/-^* mutants

Adult (>30 dpf) zebrafish arising from heterozygous *irf8* mutant incrosses were anaesthetised in 40 μg/ml tricaine methanesulfonate and a lobe of caudal fin removed by scalpel. After clipping, fins were digested to extract DNA using 10mg/ml Prot K, incubated at 65oC for 1 hour. This incubation ends with 15 minutes at 95°C to denature the Proteinase K. A section of irf8 flanking the mutation locus was then amplified from the extracted DNA by PCR using Forward -ACATAAGGCGTAGAGATTGGACG and Reverse -GAAACATAGTGCGGTCCTCATCC primers and REDTaq® ReadyMix™ PCR Reaction Mix. The PCR product was then digested for 30 minutes at 37 °C using AVA1 restriction enzyme (New England Bioscience) and the product run on a 2% agarose gel. WT = AvaI digest site is present = PCR product is cleaved to give two bands with sizes of approximately 200 and 100 bp. irf8 ^-/-^ = AvaI digest site is absent due to mutation = PCR product is not cut. A single band is observed with a size of 286 bp. irf8 ^+/-^ = Three bands as above.

### Microinjection recombinant proteins and intravital stains

Microinjections were performed on larvae at 72 hpf using a Narishige IM-300 Microinjector and pulled thin wall glass capillaries (Harvard Apparatus), administered under anaesthesia by intravenous microinjection through the cardiac sinus venosus (SV) that drains the common cardinal vein (CCV). An injection volume of 1 nL was used for all intravenous injections to minimise disruption to blood volume.

For propidium iodide intravital staining, 1nL 100μg/ml propidium iodide in DPBS was injected immediately following injury at 0.5 hpi. Larvae were then immediately imaged by heartbeat-synchronised light-sheet microscopy at 1 hpi. Injection of recombinant zfIFN-γ-rel (IFN-1.1) (Kingfisher Bioscience) was administered as a single 1nL 132nM dose at 72 hpf. Lyophilised IFN-γ-rel was reconstituted in PBS + 0.1% BSA (carrier protein) and PBS + 0.1% BSA was used as the vehicle control solution. Injections of recombinant zfVEGFaa (Kingfisher Bioscience) were administered as single 1nL 0.25 ug/ul doses at 72 hpf (protein reconstituted as above).

### Histological staining

To detect cell death at the injured ventricle, whole-mount larval TUNEL staining was performed. Larvae were fixed in 4% PFA for 30 minutes and transferred to 1:10 dilution of PBS. Larvae were subsequently digested in 1 μg/ml Proteinase K for 1 hour. Larvae were re-fixed in 4% PFA for 20 minutes and subsequently washed in PBT. TUNEL staining was performed using ApopTag Red In situ kit (MilliporeSigma) to label apoptotic cells, as described previously^26^. Stained hearts were imaged using LSFM.

EdU staining was performed by incubating larvae in 1 mM EdU (5-ethynyl-2′- deoxyuridine) (Abcam) in 1 % DMSO (Sigma Aldrich) in conditioned water + 0.003% phenylthiourea (Thermo Fisher Scientific) for 24 hours beginning either at 0 hpi or 24 hpi depending on the experiment. Larvae were incubated at 28.5°C in the dark. Larvae were then fixed for 2 hours at room temperature in 4% PFA, permeabilised in permeabilisation solution (PBS-Triton-X 0.1% + 1% Tween + 1% DMSO) and pericardium punctured using a glass microinjection needle (further improving permeability). Larvae were then washed twice in PBS-3% BSA and incubated for 2 hours at room temperature in CLICK reaction mixture from Click-iT™ EdU Imaging Kit with Alexa Fluor™ 594 (Invitrogen) made according to manufacturers’ instructions. Larvae were finally washed once in PBS-3%BSA and twice in PBS-0.1% tween and imaged by LSFM.

### Heart lesion size quantification

Larval hearts expressing the transgene *myl7:GFP* were imaged by heartbeat-synchronised light-sheet imaging as described above. Exposure was kept consistent at 10ms, along with z slice spacing (1 µm), and heart contraction phase was locked to mid diastole for all larvae. Z stacks were surface rendered in IMARIS (Bitplane) based on absolute intensity, and software-suggested segmentation and rendering parameters. Lesion area, visualised as a render-free hole in the myocardium, was then traced around manually and lesion area quantified in FIJI (National Institutes of Health)^74^.

### Ventricular ejection fraction analysis

Larval hearts of *Tg(myl7:GFP)* larvae were imaged at 80 fps in brightfield using a Leica M205 FA epifluorescence stereomicroscope, to capture when the ventricle was in diastole and systole. The ventricular area in diastole and systole was measured manually in FIJI and ventricular ejection fraction calculated using the formula 100 X [(Diastolic Area – Systolic Area)/Diastolic Area]^25^. Ventricular ejection fraction by area was then converted to ejection fraction by volume using the formula ‘Ejection fraction by area x 2.33 = Ejection fraction by volume’ derived in Supplementary Figure 4. Over the small range of ejection fractions that occur in larval hearts, the relationship can be considered to approximate to a linear one.

### Quantification of cell number by image analysis

To quantify the number of cardiomyocytes in *Tg(myl7:h2b-GFP)* and *Tg(myl7:nlsDsRed)* larval hearts, z stacks of hearts acquired by LSFM were imported into FIJI and nuclei counted using the plugin Trackmate. Briefly, key segmentation parameters ‘Estimated blob diameter’=5.5, ‘Threshold’=0.9 were taken as a starting point, and optimised manually per experiment until all nuclei are counted successfully. The heart atrium is excluded manually by x coordinate filtering and ventricular cardiomyocytes are then automatically counted by the plug in.

In order to automatically quantify the percentage of EdU+ ventricular cardiomyocytes in *Tg(myl7:h2b-GFP)* larval hearts, a custom FIJI macro was written to exclude non-cardiomyocyte EdU signal. This is necessary as cardiomyocytes have a much lower turnover rate than surrounding cells in the pericardium, endocardium and blood and so represent a minority of EdU+ cells. Briefly, the Bersen segmentation method was used to mask areas of GFP fluorescence per z slice and these masks subsequently applied as a crop RoI to EdU signal in the 641 nm colour channel of RGB images. Slices were then reassembled and merged into maximum intensity projections, where the FIJI^74^ Trackmate plugin was used to count both the total number of GFP+ cardiomyocyte nuclei and EdU+ cardiomyocyte nuclei. This quantification then allowed the percentage of EdU+ cardiomycytes to be calculated in an unbiased way per larval heart.

### Quantification of notch signalling by image analysis

In order to objectively identify whether the hearts from *Tg(Tp1:venus-PEST)* larvae possessed venus signal in the endocardium above that of background, and were therefore ‘notch+’, the following approach was used. Treatment groups were blinded to the analyser, and z stacks opened in FIJI. The automatic brightness and contrast function was used to objectively enhance the signal in the heart, and the clear interface between the granular autofluorescence of the chamber blood and the smooth autofluorescence of the myocardium searched for venus expression. The distinctive morphology and location of endothelium allowed for unambiguous identification of venus+ status.

### Pharmacological inhibition of larval signalling

To inhibit VEGF signalling, larvae were bathed in pan-VEGFR antagonist AV951/Tivozanib (Stratech Scientific) 0-48 hpi. AV951 was dissolved in 0.1% DMSO in conditioned water + 0.003% phenylthiourea to make a 10 nM solution, with just 0.1% DMSO in conditioned water + 0.003% phenylthiourea becoming the vehicle control. In order to pulse larvae with EdU, the original solution was replaced fresh solution, with the addition of 1mM EdU at 1% DMSO.

To inhibit notch signalling, larvae were bathed in gamma secretase inhibitor DAPT (Cambridge Bioscience) 0-48 hpi. DAPT was dissolved in 0.2% DMSO in conditioned water + 0.003% phenylthiourea to make a 100µM solution, with just 0.2% DMSO in conditioned water + 0.003% phenylthiourea becoming the vehicle control. Note, DAPT must be dissolved in DMSO prior to the addition of water to prevent precipitation.

In order to inhibit neuregulin-ERBB signalling, the ErBB2 antagonist AG1478 was used. Larvae were bathed in 1.75 µM AG1478 (Cambridge Bioscience) dissolved in 0.25% DMSO in conditioned water + 0.003% phenylthiourea over 0-48 hpi.

### Extraction of larval hearts and RNA extraction

Following laser injury at 72 hpf *Tg(myl7:gal4::GFP;UAS:mRFP)* larvae were incubated at 28.5°C in conditioned media/water + 0.1% methylene blue (w/v) + 0.003% phenylthiourea. At 48 hpi uninjured and injured larvae were given an overdose of tricaine at 400 μg/ml, following which hearts were extracted. We adapted the protocol of Burns and MacRae^75^ to increase the yield of heart retrieval from ∼50% to ∼70%. Briefly, ∼30 larvae were placed in 2mL eppendorf tubes, the conditioned water drained and replaced with ice cold Leibovitz’s L-15 Medium supplemented with 10% FCS. A 19-gauge needle coupled to a 5mL syringe was used to shear the larvae by aspiration and therefore dissociate hearts from the rest of the larva. The lysate was then inspected by epifluorescence microscopy and mRFP+ hearts and collected to be kept on ice. Hearts were then digested at for 10 minutes at 4°C in protease solution (5 mM CaCl2,10 mg/ml B. Licheniformis protease, 125 U/mL DNase I in 1x PBS) with occasional aspiration to aid digestion, RNA was then extracted using a RNeasy Plus Micro Kit (Qiagen) following direct lysis with RLT lysis buffer according to manufacturer’s instructions. RNA concentration was measured by Qubit and integrity by Bioanalyser. RIN score for all samples ranged between 9.6-10.

### RNAseq analysis

RNA was sequenced by Genewiz, Leipzig, Germany using Illumina NovaSeq, PE 2×150. Genewiz also used deseq2 package in R to evaluate sequencing quality, trim reads, map to the *Danio rerio* genome and generate gene counts/hits. Sequence reads were trimmed using Trimmomatic v.0.36. The trimmed reads were mapped to the *Danio rerio* GRCz10.89 reference genome available on ENSEMBL using the STAR aligner v.2.5.2b. Unique gene hit counts were calculated by using featureCounts from the Subread package v.1.5.2. Only unique reads that fell within exon regions were counted. The Wald test was used to generate p-values and log2 fold changes. A gene ontology analysis was performed on the statistically significant set of genes by implementing the software GeneSCF v.1.1-p2. The zfin GO list was used to cluster the set of genes based on their biological processes and determine their statistical significance. The volcano plot was generated by a custom R script and heatmap constructed using the pHeatmap package. For the heatmap z scaled log2(Reads) were clustered via Pearson correlation and clusters thresholded based on the resulting dendrogram. The heatmap was generated using the pHeatmap function in R.

### Statistics

Graphs and statistics were curated in GraphPad Prism 9.1 software (GraphPad Software). Data were analysed by student *t*-test, one-way ANOVA or two-way ANOVA followed by an appropriate multiple comparison *post hoc* test. All statistical tests, *p*-values and *n* numbers used are given in figure legends, p<0.05 was deemed significant in all experiments.

## Acknowledgments

This work was funded by a British Heart Foundation (BHF) CoRE award (RE/13/3/30183), Medical Research Scotland studentship (PhD-1049-2016), NC3R studentship (NC/P002196/1), BHF New Horizons grant (NH/14/2/31074), and a Medical Research Council UK award (MR/K013386/1). Bioinformatics and RNAseq performed by Genewiz, Leipzig, Germany. We also acknowledge Amelia Edmondson-Stait, University of Edinburgh, for her advice and input on RNAseq analysis.

## Author Contributions

FAB conceived of and designed the study. FAB, AK and GM carried out all experiments. Image analysis was performed by FB and AK. LSFM-related technical contributions were provided by JMT. FAB wrote the manuscript. KRS, MEMO, EGS and MB provided expertise regarding all RNA work. KRS helped optimise larval heart extraction and RNA extraction. JMT, CST, MEMO, JJM, GM, MB, AGR, and MAD edited the manuscript. MAD, AGR, and CST supervised the study. All authors contributed to the article and approved the submitted version.

## Supplementary figures

**Supplementary figure 1:**
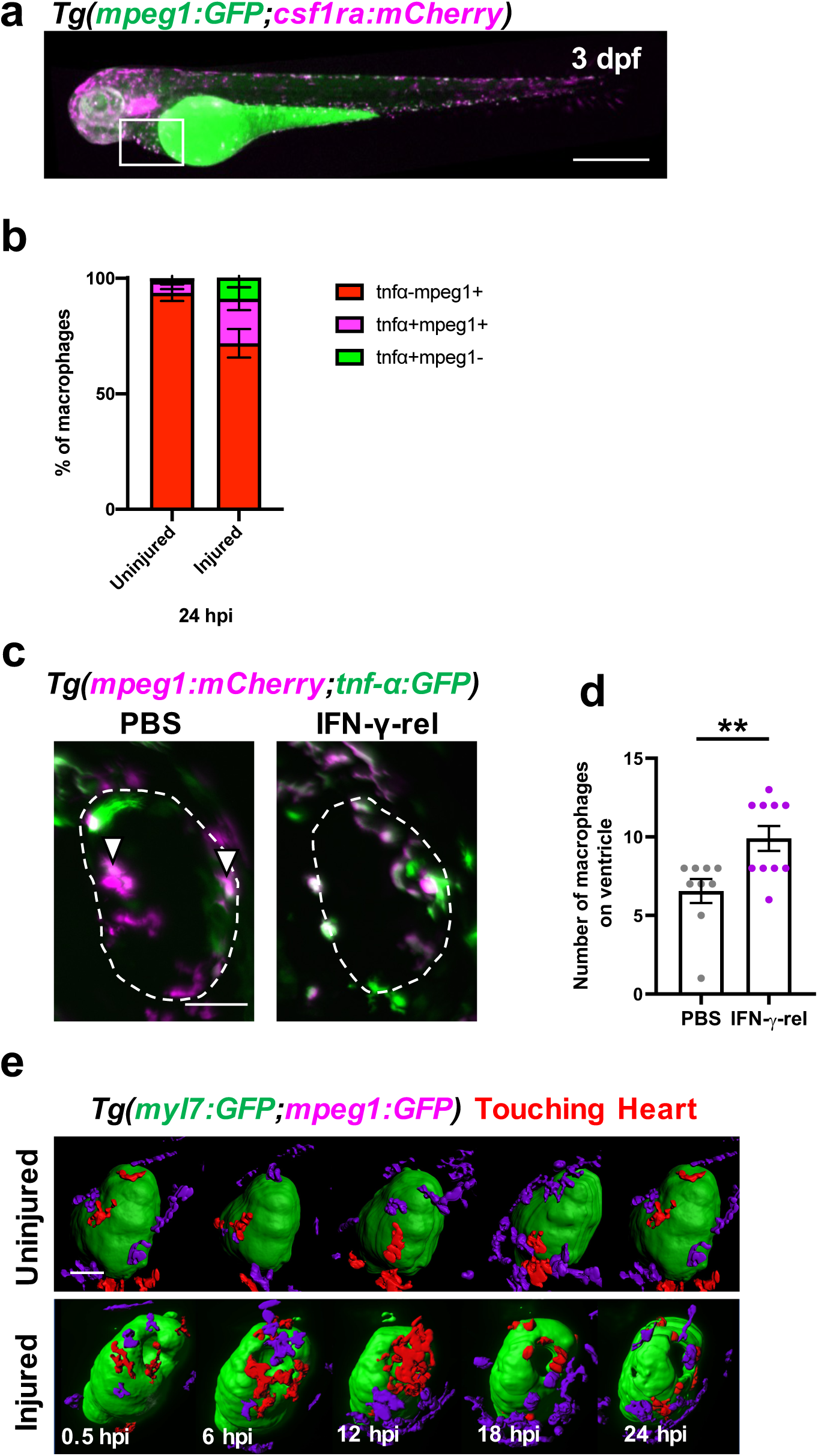
Cardiac macrophage phenotype in larval zebrafish is plastic and can be polarised to tnfa+ by IFN-γ-rel. (a) Representative epifluorescence image of a 3 dpf *Tg(mpeg1:GFP;csf1ra:NfsB-mCherry)* abbreviated to *Tg(mpeg1:GFP;csf1ra:mCherry)* in the figure, showing an anterior-posterior polarity in macrophage expression of csf1ra (higher proportion of anterior macrophages were csf1ra+). White box = indicated pericardial area. Scale bar = 500μm. (b) Quantification of the proportion of macrophages that are tnfa-mpeg1+, tnfa+mpeg1+ and tnfa+mpeg1- on hearts in uninjured and injured larvae at 24 hpi. (c) Representative images of hearts from *Tg(mpeg1:mCherry;tnfa:GFP)* larvae at 24 hpi injected with PBS or IFN-γ-rel. White dashed line = outline of the ventricle; and white arrowheads = tnfa+mpeg1+ macrophages. Scale bar = 50μm. (d) Quantification of the number of macrophages on the injured ventricle at 24 hpi after injection at 0 hpi with PBS or IFN-γ-rel. (e) Time-lapse timepoints of *Tg(myl7:GFP;mpeg1:mCherry)* hearts acquired by heartbeat-synchronised LSFM, surface rendered and colour-coded to show myocardium in green, macrophages on the heart in red and macrophages elsewhere in purple. Macrophages can be seen to change from stellate to rounded over time following injury. Scale bar = 50μm.

**Supplementary figure 2:**
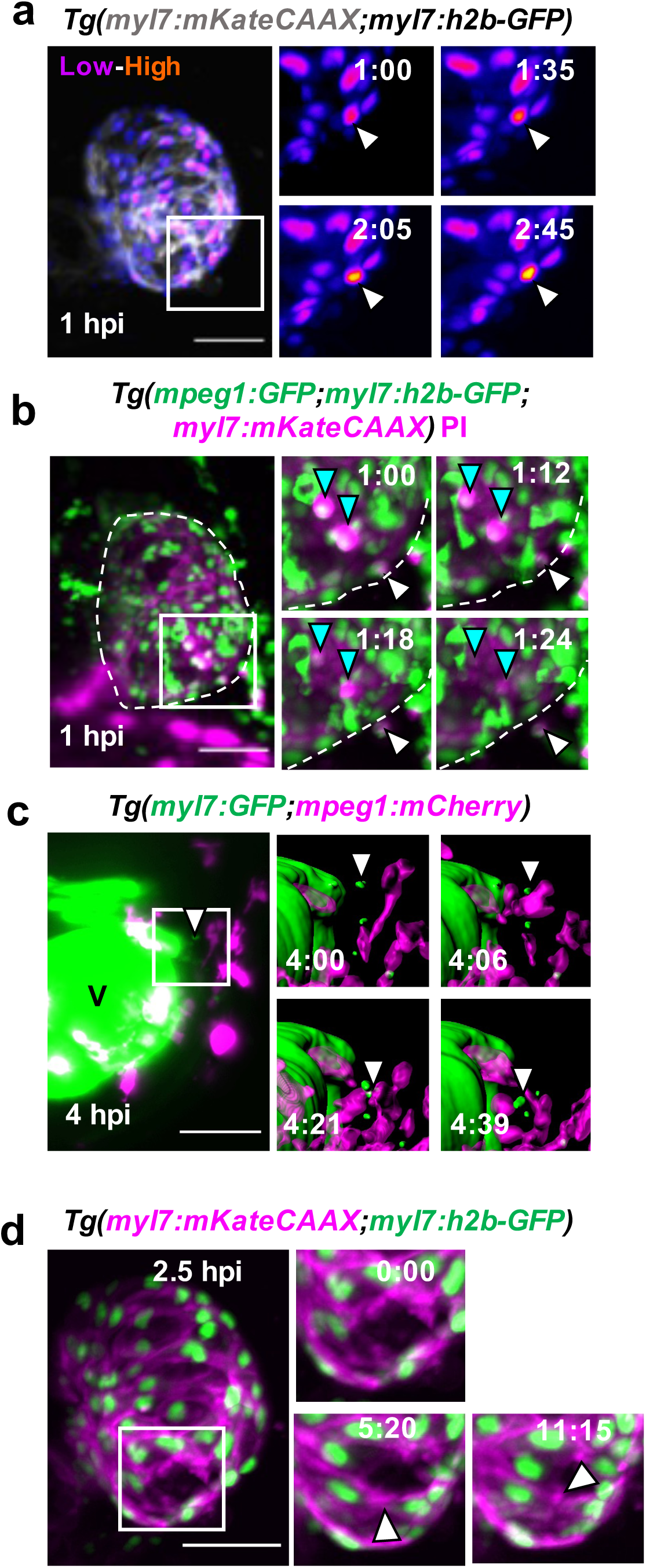
Heartbeat-synchronised lightsheet-acquired time-lapse stills. (a) Time-lapse stills of injured *Tg(myl7:h2b-GFP;myl7:mKateCAAX)* ventricles imaged from 1 hpi. GFP intensity show by heat LUT, white arrowhead = apoptotic cardiomyocyte/condensing nucleus, white box = zoom panel. (b) Time-lapse stills of injured *Tg(myl7:h2b-GFP;myl7:mKateCAAX;mpeg1:GFP)* ventricles imaged from 1 hpi by heart-synchronised light-sheet imaging. Round GFP^low^ = cardiomyocyte nuclei and stellate GFP^high^ =macrophages. Cyan arrowheads = Necrotic cardiomyocyte nuclei and white arrowheads = expelled necrotic cardiomyocyte, white box = zoom panel. (c) Time-lapse stills of an injured *Tg(myl7:GFP;mpeg1:mCherry)* ventricle from 4 hpi where the full size panel has high gain in the GFP channel to highlight GFP^low^ myocardial debris and zoom panels (area indicated by white box) are surface rendered to highlight removal of myocardium (green) by macrophages (magenta). V = high gain ventricle, white arrowhead = myocardial debris. (d) Time-lapse stills of an injured *Tg(myl7:mKateCAAX;myl7:h2b-GFP)* ventricle from 2.5 hpi. White box = zoom panel, white arrowheads = cell-cell junctions

**Supplementary figure 3:**
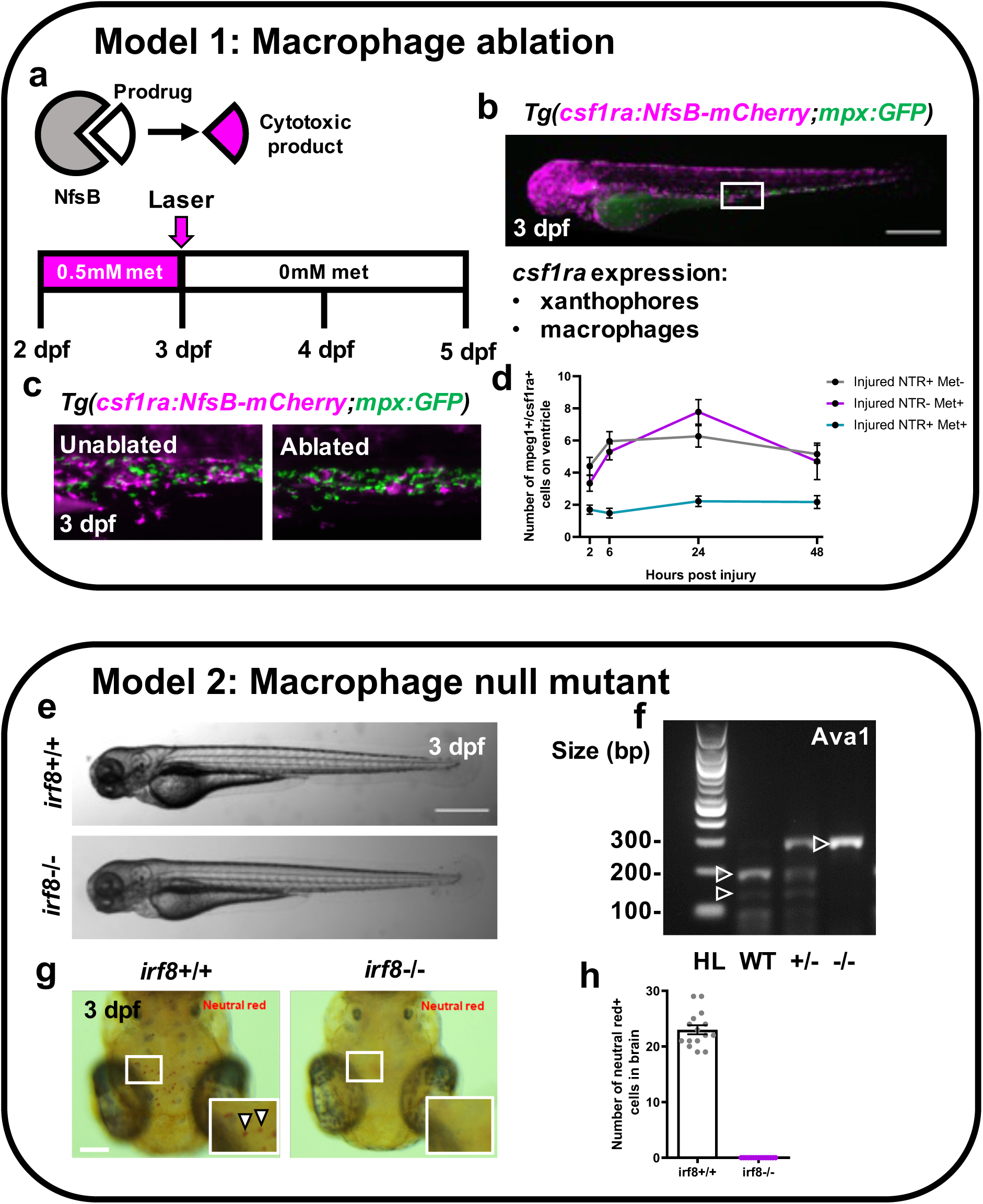
Macrophages can be pharmacologically ablated or developmentally blocked genetically. (a) Schematic illustrating how nitroreductase enzyme ‘NfsB’ catabolises prodrug ‘metronidazole’ to form a cytotoxic biproduct. (b) Representative epifluorescence image of a *Tg(csf1ra:NfsB-mCherry;mpx:GFP*) 3 dpf larva (abbreviated to *Tg(csf1ra:mCherry;mpx:GFP)* in all panels), white box = caudal haematopoietic tissue, magenta = macrophages and green = neutrophils (CHT) (c) Representative images of ablated and unablated macrophages in the CHT, size and location indicated in (b)) in *Tg(csf1ra:mCherry;mpx:GFP)* 3dpf larvae. Macrophages are ablated and only apoptotic bodies remain but not neutrophils are unaffected. (d) Quantification of macrophages at standard timepoints, marked by either mpeg1 or csfr1a on the injured ventricle in each of the NTR=metronidazole ablation model’s treatment groups NTR+Met-, NTR-Met+ and NTR+Met+. Macrophage ablation can be seen to abolish the macrophage response (e) Representative brightfield images of *irf8^+/+^* and *irf8^-/-^* larvae at 3 dpf. (f) Representative 1% agarose gel displaying Ava1 restriction digest band pattern for WT, *irf8* heterozygous and homozygous mutants. (g) Representative dorsal view brightfield image of 3 dpf larval heads stained with neutral red vital dye with white zoom panel highlighting stained macrophages (microglia) (red) in *irf8+/+* but not *irf8^-/-^* larvae. (h) Quantification of the number of neutral red positive stained cells (macrophages/microglia) in larval brains of *irf8^+/+^* and *irf8^-/-^* at 3 dpf showing *irf8^-/-^* larvae to be macrophage-null. Scale bar = 500μm (b & e), 100μm (g).

**Supplementary figure 4:**
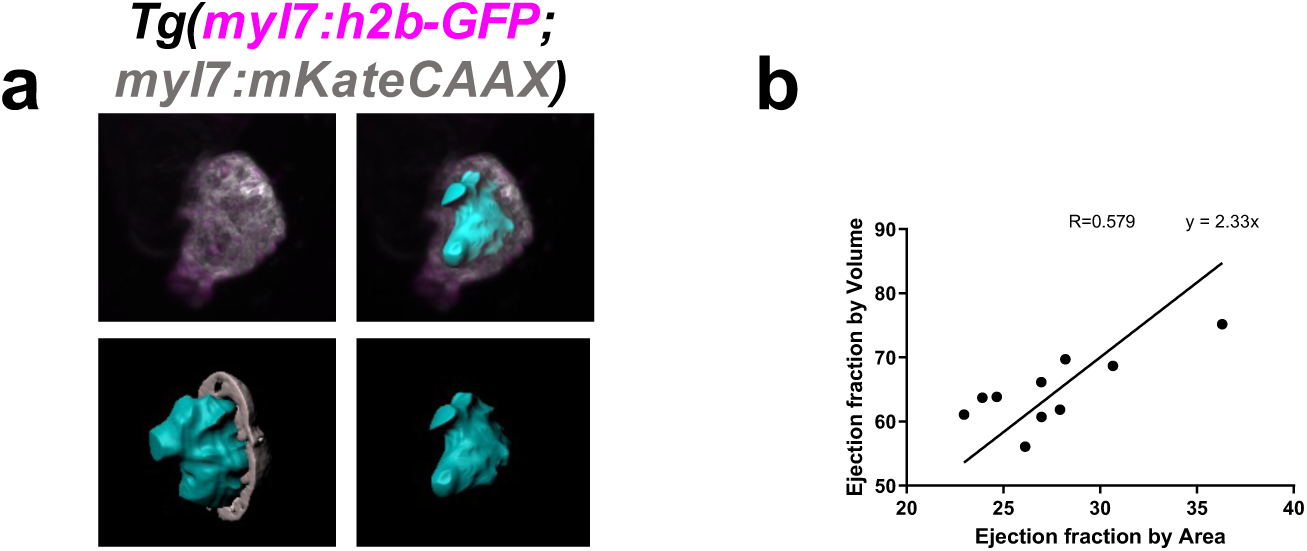
Ejection fraction by area is proportional to ejection fraction by volume. (a) Representative IMARIS-generated image showing a rendered ventricular myocardium (grey render), rendered chamber volume (cyan) and MIP of 3D heartbeat-synchronised LSFM scan of a 3 dpf heart (ventricle) in diastole. Image acquired from a *Tg(myl7:h2b-GFP;myl7:mKateCAAX)* larva. (b) Quantification of ejection fraction by area (calculated from diastolic and systolic lateral brightfield images) and by volume (calculated from surface renders of luminal volumes in diastole and systole) for n=10 fish.

**Supplementary figure 5:**
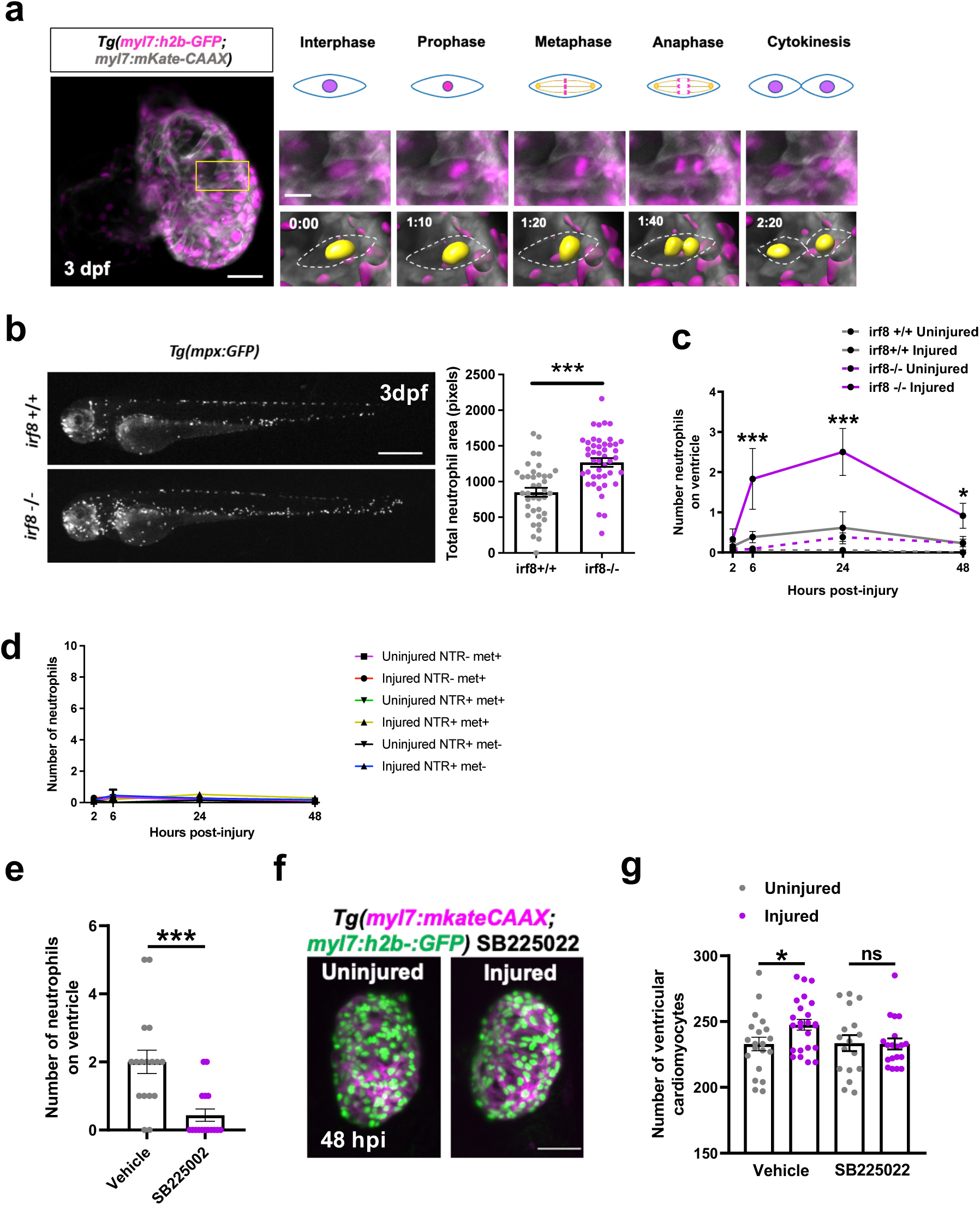
*irf8^-/-^* larvae have a larger neutrophil response to cardiac injury than *irf8^+/+^*. (a) Representative timepoint images from heartbeat-synchronised LSFM time-lapse of a laser-injured 3 dpf *Tg(myl7:h2b-GFP;myl7:mKateCAAX)* larva showing an example of each phase of complete cell division of a single cardiomyocyte, typical of larval hearts. Yellow box = zoom panel; left scale bar = 30 μm; right scale bar = 10 μm. Timestamps post-injury indicated in figure. (b) Representative whole larva epifluorescence image of *irf8^-/-^* and *irf8^+/+^ Tg(mpx:GFP)* larvae showing *irf8^-/-^* to have greater global neutrophil numbers (scale bar = 500 μm), quantified in the graph (right). \*\*\**p≤0.001.* t test, n=39-46. (c) Quantification of neutrophil numbers at the ventricle in uninjured and injured *irf8^+/+^* and *irf8^-/-^* larvae at the standard laser-injury model timepoints, showing *irf8^-/-^* larvae to have a significantly greater neutrophil response. n=17-25. (d) Quantification of neutrophil numbers at the ventricle in uninjured and injured NTR-met+, NTR+met+ and NTR+met- larvae at the standard laser-injury model timepoints. All metronidazole-nitroreductase treatment groups to have a minimal neutrophil response and therefore no neutrophil compensation in the macrophage ablated group NTR+met+, n=17-24. (e) Quantification of the number of recruited neutrophils at the injured ventricle in at 24 hpi in *Tg(myl7:h2b- GFP;myl7:mKateCAAX)* larvae bathed in vehicle or SB225002 from -2 to +24 hpi showing SB225002 to significantly reduce neutrophil number, n=17. (f) Representative light-sheet acquired images of uninjured and injured *irf8^-/-^ Tg(myl7:h2b- GFP;myl7:mKateCAAX)* ventricles at 48 hpi following treatment with SB225002 from -2 to +24 hpi, scale bar = 50 μm. (g) Quantification of ventricular cardiomyocyte number in uninjured and injured *irf8^-/-^ Tg(myl7:h2b-GFP;myl7:mKateCAAX)* ventricles at 48 hpi following treatment with vehicle or SB225002 -2 to 24 hpi, n=17-20, \**p≤0.05* t test

**Supplementary figure 6:**
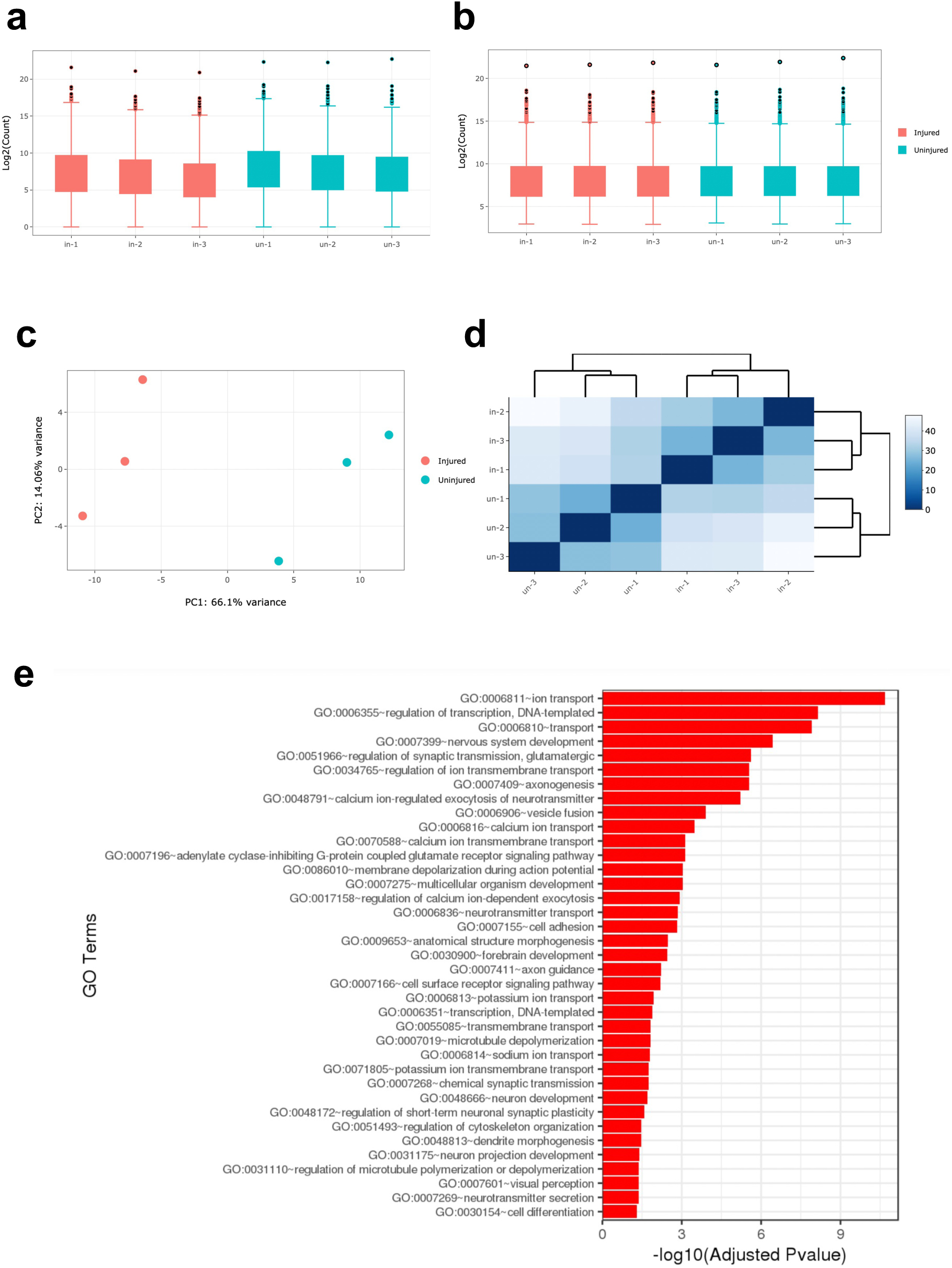
Bulk RNAseq analysis of uninjured and injured larval hearts. (a) Box plot illustrating the distribution of reads before (a) and after normalisation (b) Principal component analysis of samples, illustrating relative intragroup sample similarity. (c) Distance matrix illustrating pairwise sample similarity. (e) Plot showing gene ontology terms that were significantly enriched by Fishers exact test for significantly (padj<0.05) differentially expressed genes between uninjured and injured hearts at 48 hpi.

**Supplementary figure 7:**
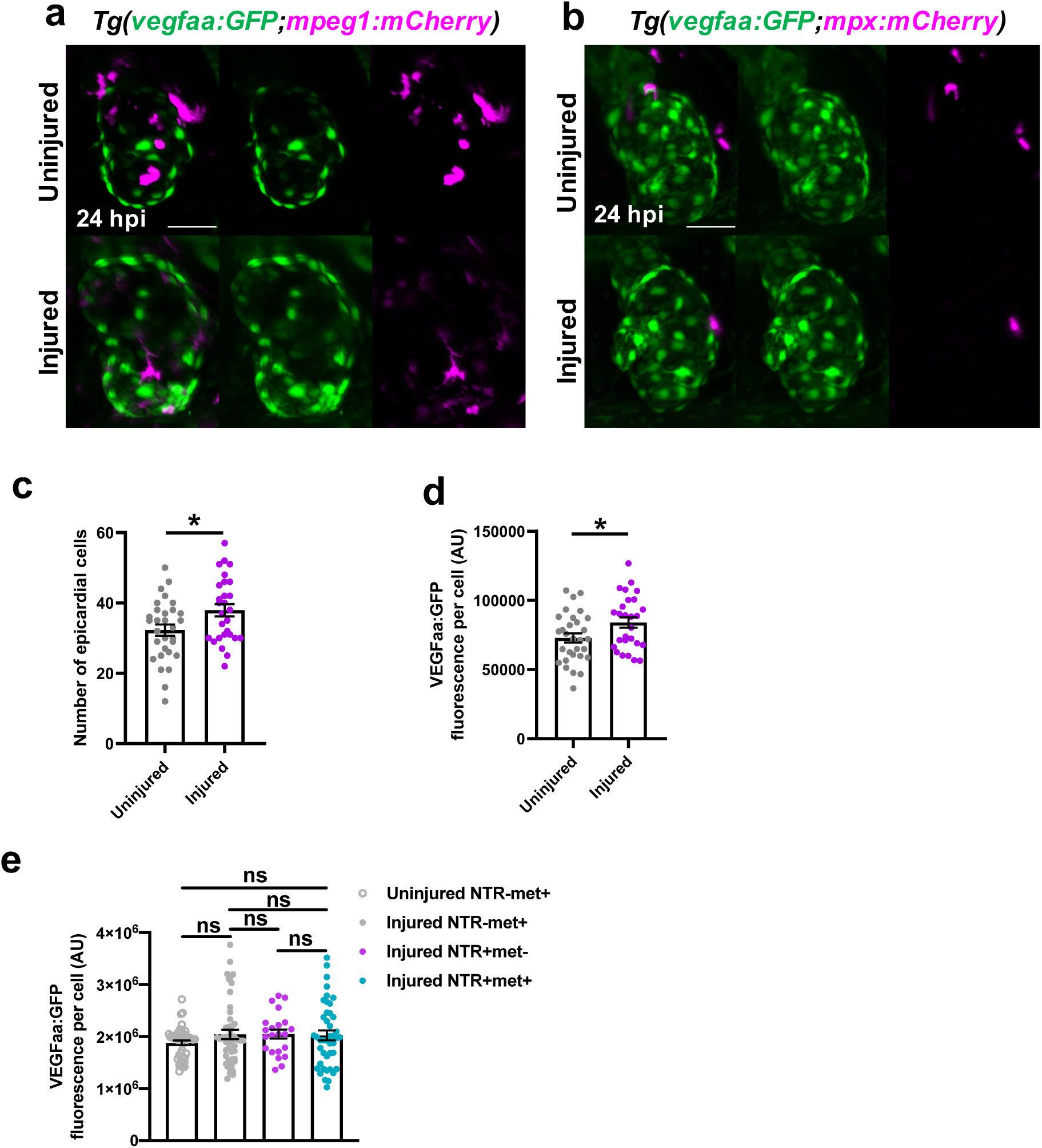
*vegfaa:GFP* expression does not colocalize with macrophages or neutrophils following larval heart injury. (a) Representative LSFM image of an injured *Tg(vegfaa:GFP;mpeg1:mCherry)* heart 24 hpi showing *vegfaa:GFP* expression only in the epicardium and not in macrophages, scale bar = 100μm. (b) Representative LSFM image of an injured *Tg(vegfaa:GFP;mpx:mCherry)* heart 24 hpi showing VEGFaa expression only in the epicardium and not in neutrophils, scale bar = 100μm. (c) Quantification of the number of epicardial cells, as marked by vegfaa:GFP, on injured ventricles at 48 hpi in uninjured and injured larvae, n=30. \**p≤0.05* t test (d) Quantification of the average vegfaa:GFP expression of epicardial cells per cell, on injured ventricles at 48 hpi in uninjured and injured larvae, n=30. \**p≤0.05* t test (e) Quantification of average vegfaa:GFP fluorescence per cell in metronidazole-nitroreductase ablation model groups at 48 hpi, n=22-44. One-way ANOVA followed by Holm-Sidak’s multiple comparisons Post-hoc test.

## Videos

Videos 1-8 can be found deposited on Dropbox via the following link:

https://www.dropbox.com/sh/xy5re3qvf4oc327/AAAVYj6SlCxZBJGKKYQLNI6Ia?dl=0

Video 1: LSFM-acquired heartbeat-synchronised time-lapse of a *Tg(csf1ra:mCherry;mpeg1:GFP)* heart showing macrophage heterogeneity following cardiac injury.

Video 2: LSFM-acquired heartbeat-synchronised time-lapse of a *Tg(mpeg1:mCherry;tnfa:GFP)* heart showing macrophage plasticity following cardiac injury.

Video 3: LSFM-acquired heartbeat-synchronised time-lapse of a *Tg(myl7:h2b-GFP;myl7:mKateCAAX)* heart following cardiac injury showing cardiomyocyte apoptosis following injury.

Video 4: LSFM-acquired heartbeat-synchronised time-lapse of a *Tg(myl7:h2b-GFP;mpeg1:GFP)* heart injected with propdium iodide showing PI+ cardiomyocyte expulsion following cardiac injury.

Video 5: LSFM-acquired heartbeat-synchronised time-lapse of a *Tg(myl7:GFP;mpeg1:mCherry)* heart, 3D surface rendered, showing removal and internalization of myocardial debris by macrophages following injury.

Video 6: LSFM-acquired heartbeat-synchronised time-lapse of a *Tg(myl7:GFP)* heart, 3D surface rendered, showing budding and bridging of wound margin myocardium following injury.

Video 7: LSFM-acquired heartbeat-synchronised time-lapse of a *Tg(myl7:h2b-GFP;myl7:mKateCAAX)* heart, showing budding and bridging of individual wound-margin cardiomyocytes following injury.

Video 8: LSFM-acquired heartbeat-synchronised time-lapse of a *Tg(myl7:h2b-GFP;myl7:mKateCAAX)* heart, showing cardiomyocyte cell division with nuclear division and cytokinesis.

